# Epithelial tubule interconnection driven by HGF-Met signaling in the kidney

**DOI:** 10.1101/2024.06.03.597185

**Authors:** Isabel López-García, Sunhee Oh, Chris Chaney, Jun Tsunezumi, Iain Drummond, Leif Oxburgh, Thomas Carroll, Denise K. Marciano

## Abstract

The formation of functional epithelial tubules is a central feature of many organ systems. Although the process of tubule formation by epithelial cells is well-studied, the way in which tubules connect with each other (i.e. anastomose) to form functional networks both *in vivo* and *in vitro* is not well understood. A key, unanswered question in the kidney is how the renal vesicles of the embryonic kidney connect with the nascent collecting ducts to form a continuous urinary system. We performed a ligand-receptor pair analysis on single cell RNA-seq data from embryonic mouse kidney tubules undergoing anastomosis to select candidates that might mediate this process *in vivo*. This analysis identified hepatocyte growth factor (HGF), which has known roles in cell proliferation, migration, and tubulogenesis, as one of several possible candidates. To test this possibility, we designed a novel assay to quantitatively examine epithelial tubule anastomosis *in vitro* using epithelial spheroids with fluorescently-tagged apical surfaces to enable direct visualization of anastomosis. This revealed that HGF is a potent inducer of tubule anastomosis. Tubule anastomosis occurs through a proliferation-independent mechanism that acts through the MAPK signaling cascade and matrix metalloproteinases (MMPs), the latter suggestive of a role in extracellular matrix turnover. Accordingly, treatment of explanted embryonic mouse kidneys with HGF and collagenase was sufficient to induce kidney tubule anastomosis. These results lay the groundwork for investigating how to promote functional interconnections between tubular epithelia, which have important clinical implications for utilizing *in vitro* grown kidney tissue in transplant medicine.

## INTRODUCTION

Tubular structures underlie the functions of many tissues—from the blood vasculature, to the digestive tract, to the nephrons of the kidney. The adult human kidney is comprised of approximately 0.5 to 1 million nephrons (1, 2), each of which consists of a filtration unit at its proximal end which then feeds into a long, segmented epithelial tubule. Throughout this tubule, the nephron filtrate undergoes significant modifications to reclaim nutrients, expel waste, and modify urine concentration. Disruptions to nephron filtration or tubular functions can lead to kidney disease (3, 4). In severe cases, individuals with impaired kidney function may require renal replacement therapy, typically via dialysis or kidney transplant. Despite their life-extending benefits, both treatments have limitations, underscoring the urgent need for alternative strategies to augment kidney function.

One potential alternative strategy is implantation of stem cell-derived kidney tissue into individuals with severely impaired kidney function. Thus far, most efforts to generate kidney tissue from pluripotent stem cells have focused on how to maximize tubule differentiation, segmentation, and cellular function (5–9), and how to support tubule vascularization (10–13). Current methods are based on promoting differentiation of stem cells by exposing them to a series of factor supplementations (5, 14–18). This method, while promising, faces barriers for use in future transplant medicine. Central to this endeavor is the creation of functional interfaces between implanted and recipient tissues to ensure concerted function. To that end, facilitating anastomosis of engineered tubules with the tubules of a recipient host remains a major hurdle.

Nature has already solved the challenge of engineering epithelial tubule interconnections. In the developing mammalian kidney, mesenchymal nephron progenitors undergo a mesenchymal-to-epithelial transition to form epithelial spheroids called renal vesicles. Adjacent to these renal vesicles are the tips of ureteric tubules, or ureteric buds, which give rise to mature collecting ducts. During nephrogenesis, the renal vesicles elongate and then anastomose with tips of the ureteric buds to form continuous tubules that then undergo lumen fusion to yield a single central lumen (19, 20).

Work from our lab has previously demonstrated that the lumenization of these anastomosed tubules occurs by de novo lumen formation as well as lumen fusion (21). Yet, what drives the process of tubule anastomosis between a developing renal vesicle and the ureteric bud remains unknown. This is in part due to the complexity of the system, as there are numerous developmental and morphogenetic processes occurring, such as differentiation and progenitor renewal (22–25), cell migration (26), matrix secretion and turnover (27–29), and concurrent mesenchymal-to-epithelial transition. In addition, the tubules themselves are surrounded by progenitor interstitial cells that play a role in cell differentiation and morphogenesis (30), although their role in tubule anastomosis is unknown. Other model systems that undergo spontaneous epithelial tubular anastomosis including the drosophila trachea (31), avian lung (32), and zebrafish kidney (33), are all similarly complex.

To assess the molecular underpinnings of epithelial tubule anastomosis and to identify candidate effectors, we performed a ligand-receptor pair analysis on single cell RNA-seq data from embryonic mouse kidney tubules undergoing anastomosis (34–37). In conjunction with these studies, we developed a novel fluorescence-based, quantitative 3D tissue culture model designed to identify cellular and molecular mechanisms leading to tubule anastomosis and lumen fusion. This assay builds upon a well-characterized three-dimensional (3-D) cell culture system that generates epithelial spheroids/tubuloids from Madin-Darby Canine Kidney (MDCK) cells (38–41) and can be used to test factors that promote tubule anastomoses in a directed fashion. One of the top candidates identified in the ligand-receptor analysis, hepatocyte growth factor (HGF), was found to be sufficient to induce tubule anastomoses in the tubuloid assay, where it does so in a proliferation-independent, MAPK-dependent manner. HGF was also found to induce anastomosis of tubules from mouse kidney explants, thus validating the *in vitro* approach.

Given the numerous clinical applications for promoting functional tubular interconnections, particularly at the intersection of organoid biology and transplantation, these findings are certain to have a direct and significant impact. Furthermore, this novel assay, combined with knowledge of candidate pathways, offers a powerful tool for rapid hypothesis testing in a variety of different settings.

## RESULTS

### A bioinformatics-based approach to uncover mechanisms underlying interconnection in vivo

During normal nephrogenesis, the tips of the ureteric bud and the renal vesicle anastomose to form a single continuous conduit, schematically represented as **Fig 1A**. Although little is known about the underlying mechanisms, we reasoned that the interconnection between these tubules might involve concerted interactions at the level of surface proteins. To identify ligand-receptor pairs that might be involved, we utilized previously published single cell RNA-seq datasets from mouse kidney during late embryogenesis (34, 35) and performed ligand-receptor analysis with the cell clusters representing the tips of the ureteric bud (UB) and the distal portion of the renal vesicle (RV). Additionally, because the UB and RV tubules are adjacent to interstitial cells that may secrete morphogens or provide other cues, we also performed ligand-receptor analysis between the UB/RV and the surrounding three nephrogenic stromal/interstitial cell populations, referred to as stromal progenitors SP1, SP2, and SP3. The clusters representing these individual cell types are shown as a UMAP plot (**Fig 1B**) and select results of the ligand-receptor analysis are represented as a ladder plot (**Fig 1C**, see **Table S1** for top 100 interactions). Amongst these results, hepatocyte growth factor (HGF) emerged as a highly ranked ligand (top 25) four times in conjunction with MET and ST14 in the SP2-UB and SP3-UB analyses. MET is a tyrosine kinase receptor that activates HGF signaling (42), while ST14 is a transmembrane serine protease that activates pro-HGF (43).

**Figure 1.**
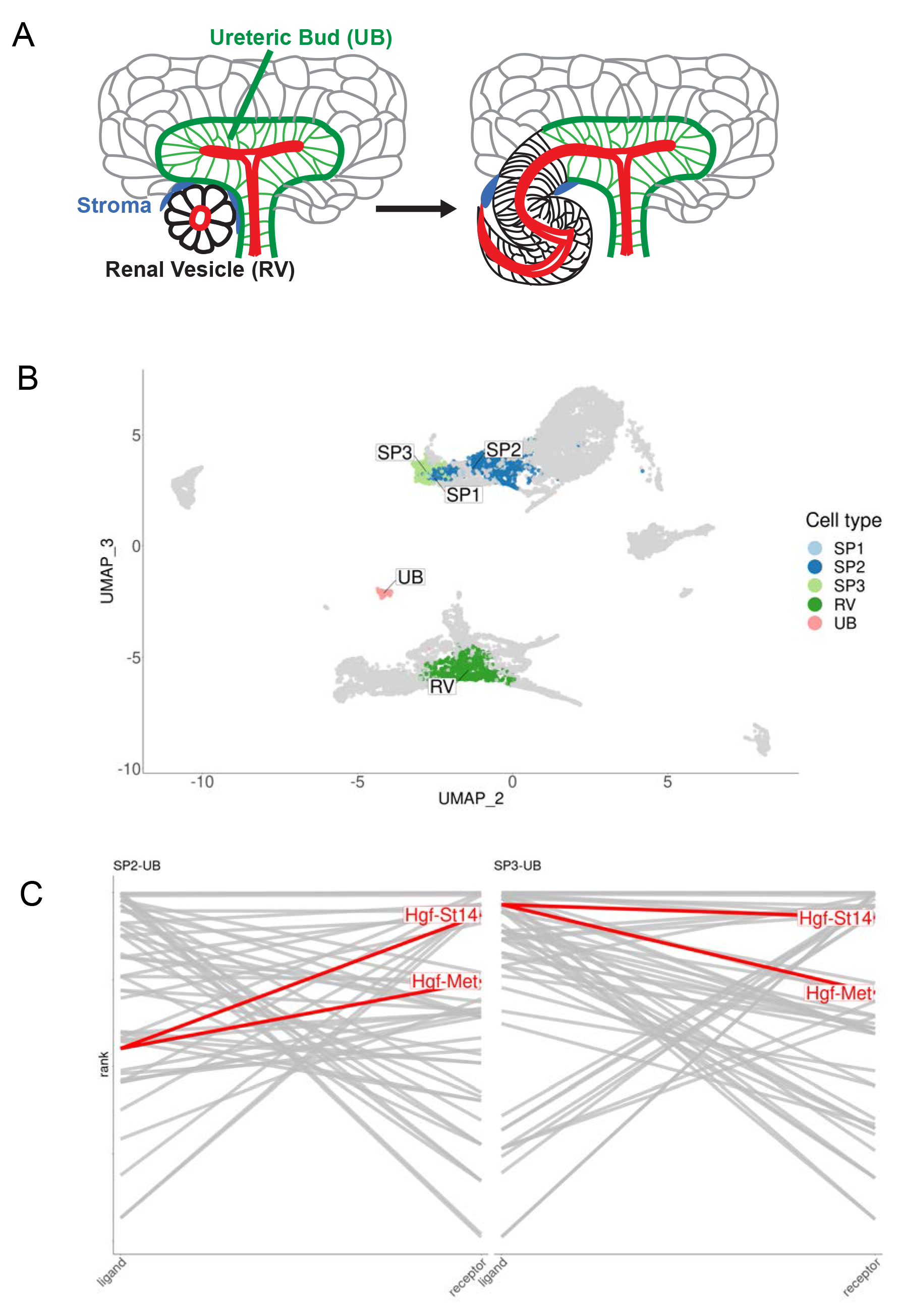
Ligand-receptor analysis identifies tubule anastomosis-promoting candidate genes. **A** Schematic of tubule anastomosis in developing mammalian kidney. The distal renal vesicle (black) connects with tips of the ureteric tubule (green). Nephrogenic stromal/interstitial cells (blue) surround the site of anastomosis. **(B)** UMAP indicating the distal RV, UB tips, and nephrogenic stromal/interstitial cell populations PS1, PS2, and PS3 in E18.5 mouse kidneys. **(C)** Ligand-receptor (L-R) ladder plots are shown for SP2-UB and SP3-UB. (SP1 contained very few cells.) For a given pair of interacting cell types, ligands are ranked in descending order by expression specificity in the source population on the left-hand side and receptors ranked in descending order of expression specificity in the target population on the right-hand side. A line segment connects the ligands to receptors of interacting pairs. The higher the “rung of the ladder”, the more specific the ligand-receptor interaction. The top 25 L-R pairs for each analysis are indicated. HGF and its two identified receptors, c-Met and St14, are highly ranked in both. See Table S1 for top 100 L-R interactions between all interacting cell types.

### MDCK spheroids anastomose over time in 3D culture

When single cell-dissociated MDCK cells are grown in Matrigel, each will proliferate and form a polarized 3D spheroid, as previously described (44, 45). We observed over time that some of the spheroids were dumbbell-shaped, suggesting spheroid anastomosis (**Fig 2A**). To examine this phenomenon more closely, we generated MDCK spheroids stably producing fluorescently-tagged podocalyxin. Although fluorescently-tagged podocalyxin (Podxl) localizes to the apical plasma membrane, through partial apical ectodomain shedding (46), these N-terminally-tagged fluorophores also highlight the lumenal space. Visualization of GFP-Podxl-tagged spheroids revealed that some spheroids “invaded” adjacent spheroids (**Fig 2Ba**). Other spheroids appeared connected but had two distinct lumens (**Fig 2Bb**), while dumbbell-shaped spheroids with two main spheroidal subdomains contained a single lumen (**Fig 2Bc**).

**Figure 2.**
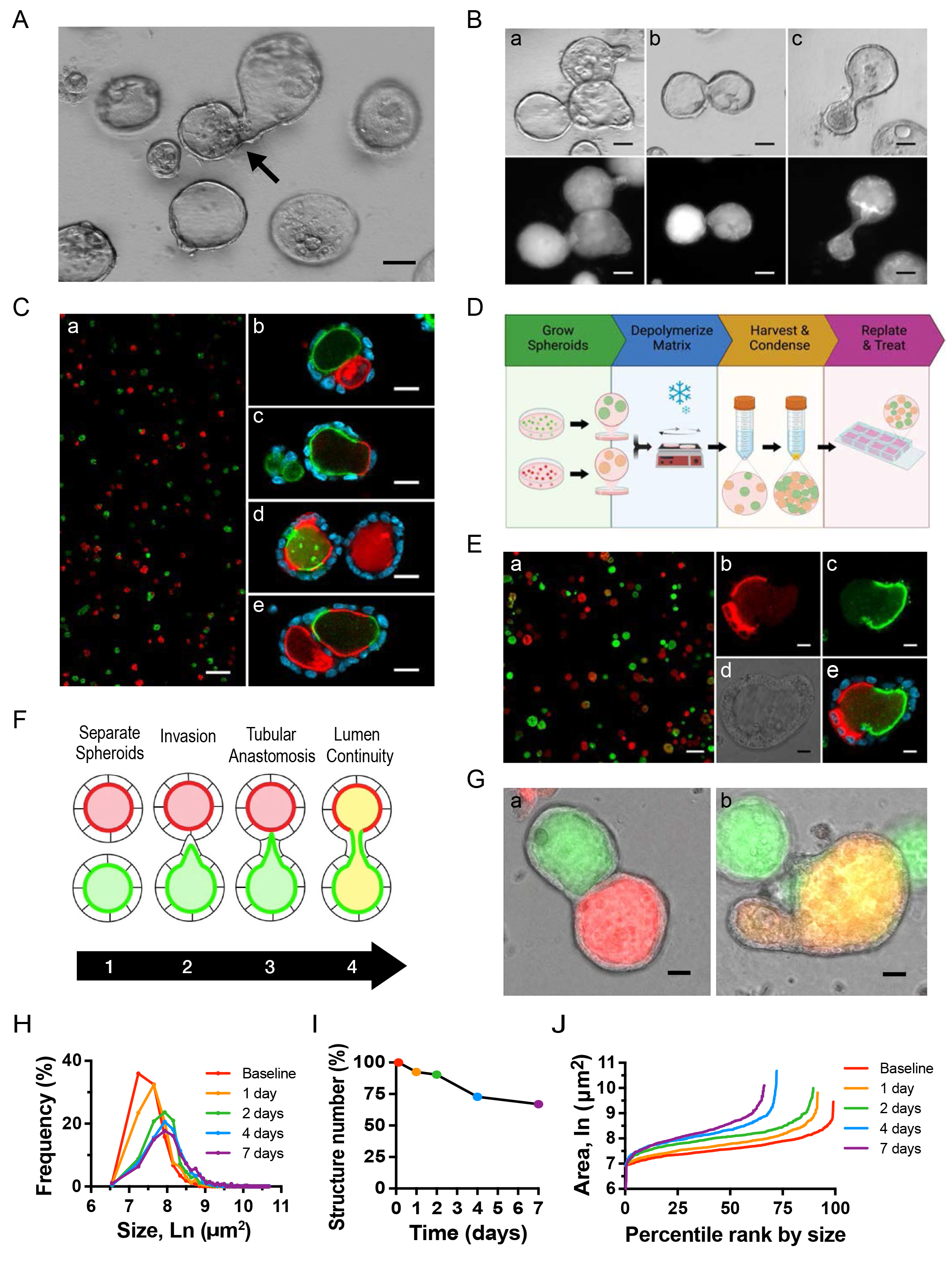
Development of a quantitative assay of spheroid interconnection. **A** Brightfield imaging of 2-wk-old spheroids depicts non-spherical structures consistent with anastomosis. **(B)** Brightfield and fluorescent images of GFP-Podxl^+^ spheroids show asymmetrical protrusions (a), adjacent spheroids with a common basal surface (b), and a dumbbell-shaped structure with a single lumen (c). Images are representative of at least five independent experiments. (A, B) Images are representative of at least five independent experiments. Scale bars = 50 µm. **(C)** Fluorescent images from experiments in which RFP-Podxl^+^ and GFP-Podxl^+^ cells were mixed prior to spheroid formation (a) show single (b), double (c), and intercalated (d,e) fluorescently- labeled lumens. **(D)** Schematic of new assay in which RFP-Podxl^+^ and GFP-Podxl^+^ spheroids are grown separately, harvested, condensed, and replated. **(E)** Fluorescent images from new assay show single or dual-labeled lumens without intercalation. High magnification fluorescent (b,c,e) and brightfield (d) images of a dual-fluorescent structure are shown. Images are representative of at least ten independent experiments. Scale bars = 200 µm (Ca, Ea), 20 µm (Cb-e), 10 µm (Eb- e) **(F)** Schematic of the stages of anastomosis. **(G-J)** Spheroid anastomosis was quantified by four measures: **(G)** Structures with dual GFP^+^ and RFP^+^ fluorescence (tubular anastomosis) shown as percentage of total structures (a). Structures with lumen spectral overlap of GFP^+^ and RFP^+^ or ‘yellow’ lumens as percentage of total structures (b). Images are representative of at least ten independent experiments. Scale bars = 20 µm. (H) Frequency distribution of structure cross-sectional area over time. (I) Structure number as percentage of baseline count. This decreases over time due to anastomoses. Spheroid size (H) and number (I) can also be represented in a single graph, as in (J).

### A novel assay for spheroid interconnection

To determine if dumbbell-shaped spheroids were two anastomosed spheroids rather than a single spheroid with unusual shape, we generated two cell lines stably producing GFP-Podxl or RFP- Podxl. Dissociated cells from both cell lines were plated at a 1:1 ratio and grown into spheroids (**Fig 2Ca**). By examining both fluorophores, heterologous connection of GFP^+^ and RFP^+^ spheroids could be detected. Some fused structures had separate GFP^+^ and RFP^+^ lumens (**Fig 2Cb**) while others had a single lumen that was double-positive (GFP^+^-RFP^+^) (**Fig 2Cc**). Both types of structures were suggestive of anastomosis, with the former in an intermediate state that had not undergone lumen fusion. However, we also observed varying degrees of intercalation between GFP^+^ and RFP^+^ apical fluorescence in some spheroids (**Fig 2Cd,e**). Thus, it was not clear if these GFP^+^-RFP^+^ structures arose from connection of appropriately polarized and lumenized spheroids or from epithelial cells that had aggregated prior to spheroid formation and lumenization.

To distinguish between these possibilities and detect only anastomosed spheroids, we created a new assay in which GFP-Podxl^+^ and RFP-Podxl^+^ spheroids were grown separately, detached from their extracellular matrix (ECM) substrate, and then re-plated together (**Fig 2D**). Briefly, spheroids were harvested and re-plated after 3 days in Matrigel culture, when spheroids had apical-basal polarity and discrete lumens (47). We optimized the assay by increasing the re- plated density of spheroids 10-fold after centrifugation and resuspension to increase the basal rate of interconnection. In this assay, a single structure containing GFP^+^ and RFP^+^ could only be the result of interconnection between separate spheroids. Figure 2E depicts co-plated spheroids (**Fig 2Ea**) and highlights an example of a structure resulting from two anastomosed spheroids (**Fig 2Eb-e**).

### Quantification of spheroid anastomosis by fluorescence- and morphology-based metrics

Using this assay, we examined the process of normal anastomosis at cellular resolution. During interconnection, two spheroids progress from invasion and tubular anastomosis to lumen fusion (**Fig 2F**). The patterns of fluorescence provide direct readouts of the extent of interconnection. The first stage shows dual GFP^+^-RFP^+^ structures (**Fig 2Ga**). The latter shows spectral overlap between GFP^+^ and RFP^+^ at coalesced lumens (**Fig 2Gb**), giving them a yellow appearance on epifluorescence imaging. In addition to the robust, fluorescence-based, binary readouts (dual versus single color; yellow lumen present versus absent), changes in spheroid size and number can be used as indirect measures of spheroid, and subsequently lumen, interconnection (**Fig 2H- J**). For example, the cross-sectional area of interconnected structures is larger than that of single spheroids that have only proliferated over the same time. Similarly, a decrease in the number of structures over time in each area suggests interconnection has occurred. Collectively, these four metrics provide strong quantitative readouts of spheroid interconnection.

### Hepatocyte growth factor promotes spheroid anastomosis

Among known tubulogens (factors promoting tubule formation in 3D cultures), HGF has a well-known role in inducing cellular protrusions and influencing tubule morphogenesis, depending on the extracellular context (48–52). Based on this, we hypothesized that HGF might also promote tubule anastomosis and selected it for further study. We added HGF to replated spheroids and observed dramatic and robust effects on tubule interconnection (**Fig 3A**). As previously described, spheroid interconnection was assessed based on four metrics: the percentage of 1) dual GFP^+^- RFP^+^ structures, 2) GFP^+^-RFP^+^ (‘yellow’) lumens, 3) spheroid size, and 4) spheroid number (**Fig 3B,C,D,E,F**). HGF promoted spheroid anastomosis in a dose-dependent manner (**Fig S1**).

**Figure 3.**
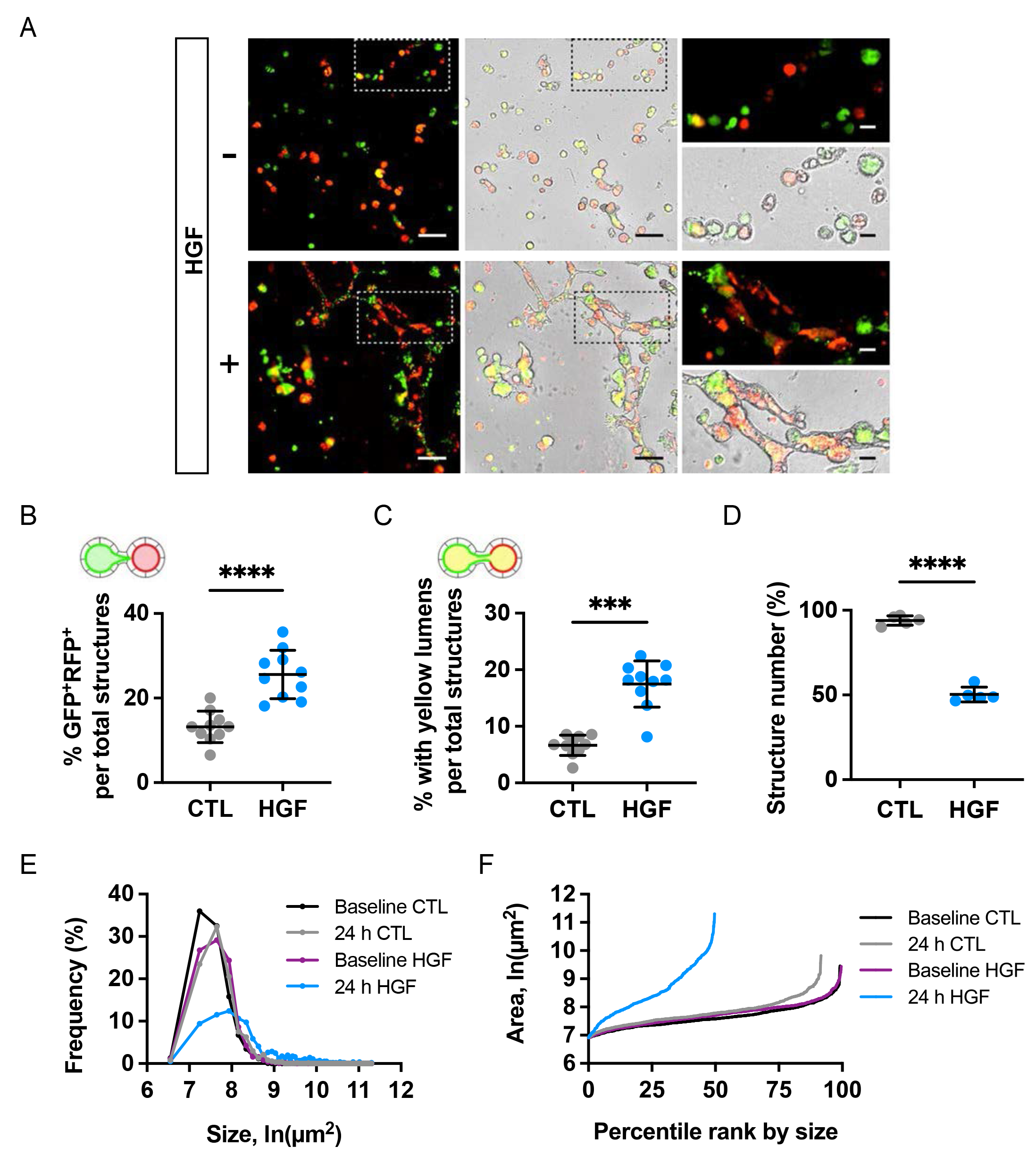
HGF promotes extensive interconnection in replated MDCK spheroids. Replated spheroids were treated with HGF (10 ng/mL) or vehicle for 4 days. Media was replenished every two days. **(A)** Representative brightfield and fluorescent images at study endpoint. Scale bars =200 µm; 50 µm (insets). Measures of interconnection were scored by blind observer. **(B)** Percentage of GFP^+^-RFP^+^ structures. **(C)** Percentage of structures with ‘yellow’ lumens. **(D)** Structure number as a percentage of baseline. **(E)** Frequency distributions of cross-sectional areas of all identified structures by treatment group. **(F)** Structure count versus size plot after 24 hours of treatment. Note reduction in structure count and increase in structure sizes, reflecting HGF-induced interconnection. Data are representative of at least ten independent experiments. Measures of interconnection and number were compared by Student’s t-test: *P*&lt;.0001****, *P*&lt;.001***.

### Proliferation is not necessary for tubular interconnection

Since a major function of HGF is to promote cellular proliferation, we tested the role of cell proliferation in tubule anastomosis using the inhibitor mitomycin C. Preliminary experiments were performed on MDCK spheroids to titrate the dose of mitomycin C that impeded cell proliferation without inducing cell death (**Fig S2**). We found that a proliferation-inhibiting dose of mitomycin C did not inhibit anastomosis by any of the four quantitative measures (**Fig 4A-D**) but decreased the size of spheroids (**Fig 4D**) and their elongation (**Fig 4A**), resulting in mostly spherical structures. This suggests that anastomosis and proliferation can be molecularly uncoupled, where proliferation, resulting in larger, irregularly shaped spheroids, may occur normally after HGF treatment, but it is not required *a priori* for epithelial fusion downstream of HGF.

**Figure 4.**
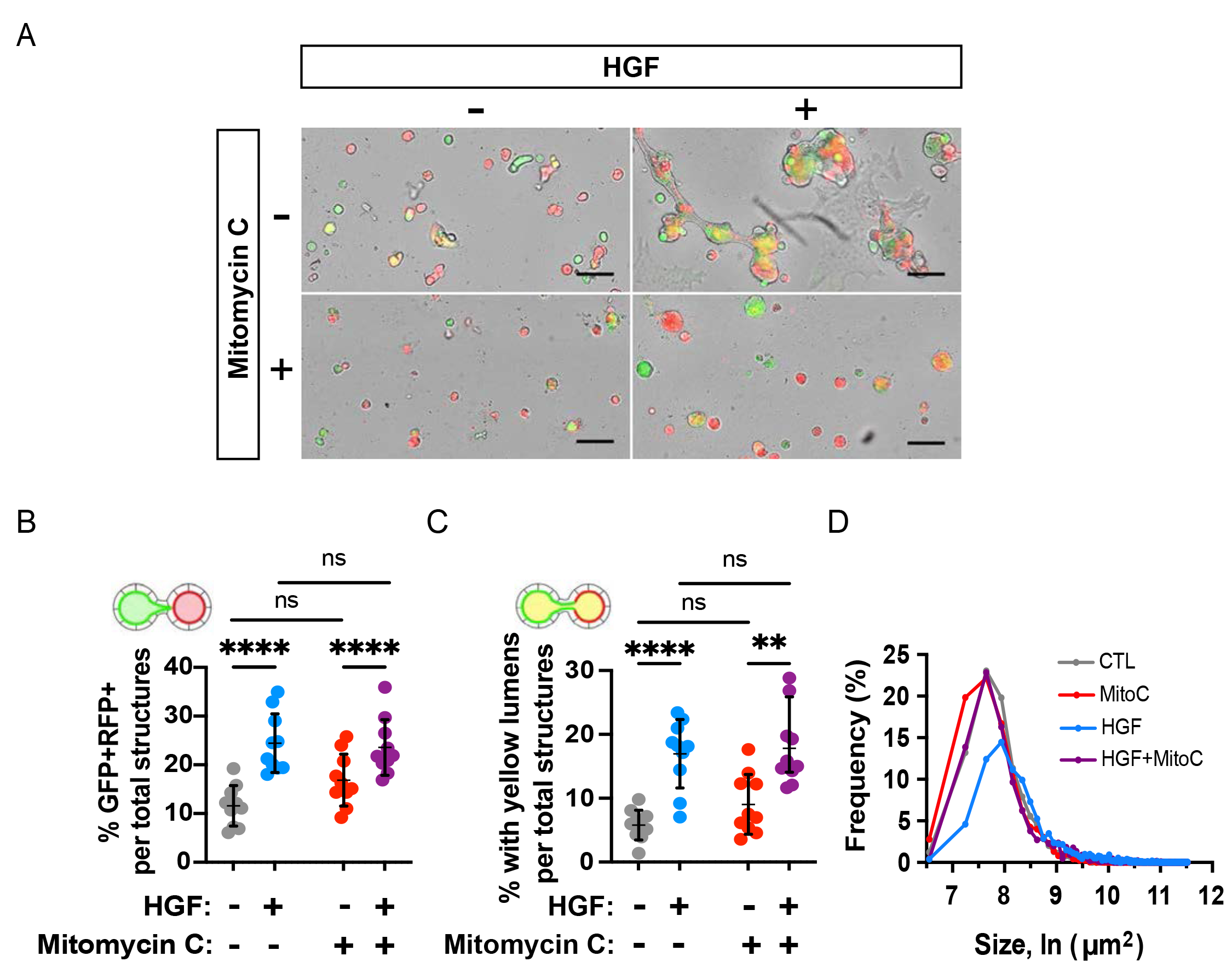
Proliferation is dispensable for HGF-induced anastomosis. Replated spheroids were treated with either HGF (10 ng/mL), an inhibitor of cell proliferation, mitomycin C (2.5 µM), or a combination, for two days. **(A)** Representative images of replated spheroids after treatment period. Scale bars = 200 µm. **(B)** Percentage of GFP^+^-RFP^+^ structures were not altered by inhibitor treatment. **(C)** Percentage of yellow lumens was unaffected by mitomycin C. **(D)** Cross-sectional area of structures at endpoint. Data shown are representative of at least four independent experiments performed in triplicate. Measures of interconnection were compared by two-way ANOVA, *P*&lt;.0001****.

### HGF activates the MEK/ERK signaling pathway to stimulate tubular interconnection

HGF triggers pleiotropic cellular responses by binding MET receptor tyrosine kinase, activating multiple downstream signaling pathways, including STAT3 and MAPK (i.e. MEK/ERK) signaling (53, 54). HGF triggers phosphorylation of the Tyr705 residue of STAT3 (55), and promotes scattering and morphogenesis of epithelial cells by activating ERK1/2 (53, 56). To determine how HGF stimulates tubular interconnection, we examined the phosphorylation status of STAT3 and ERK1/2 in replated MDCK spheroids in the presence or absence of HGF (**Fig 5A**). While phosphorylation of the Ser727 residue of STAT3 remained constant, phosphorylation of the Tyr705 residue increased between 0.5 h and 2 h after culture, regardless of the HGF treatment. This indicates that HGF does not affect the phosphorylation status of STAT3 in the replated spheroids. In contrast, the same samples had a robust increase ERK1/2 phosphorylation in the presence of HGF compared with those lacking HGF at all time points assayed (**Fig 5A**). The results show that HGF activates ERK1/2 MAP kinase in MDCK spheroids and does so within 30 minutes of treatment.

**Figure 5.**
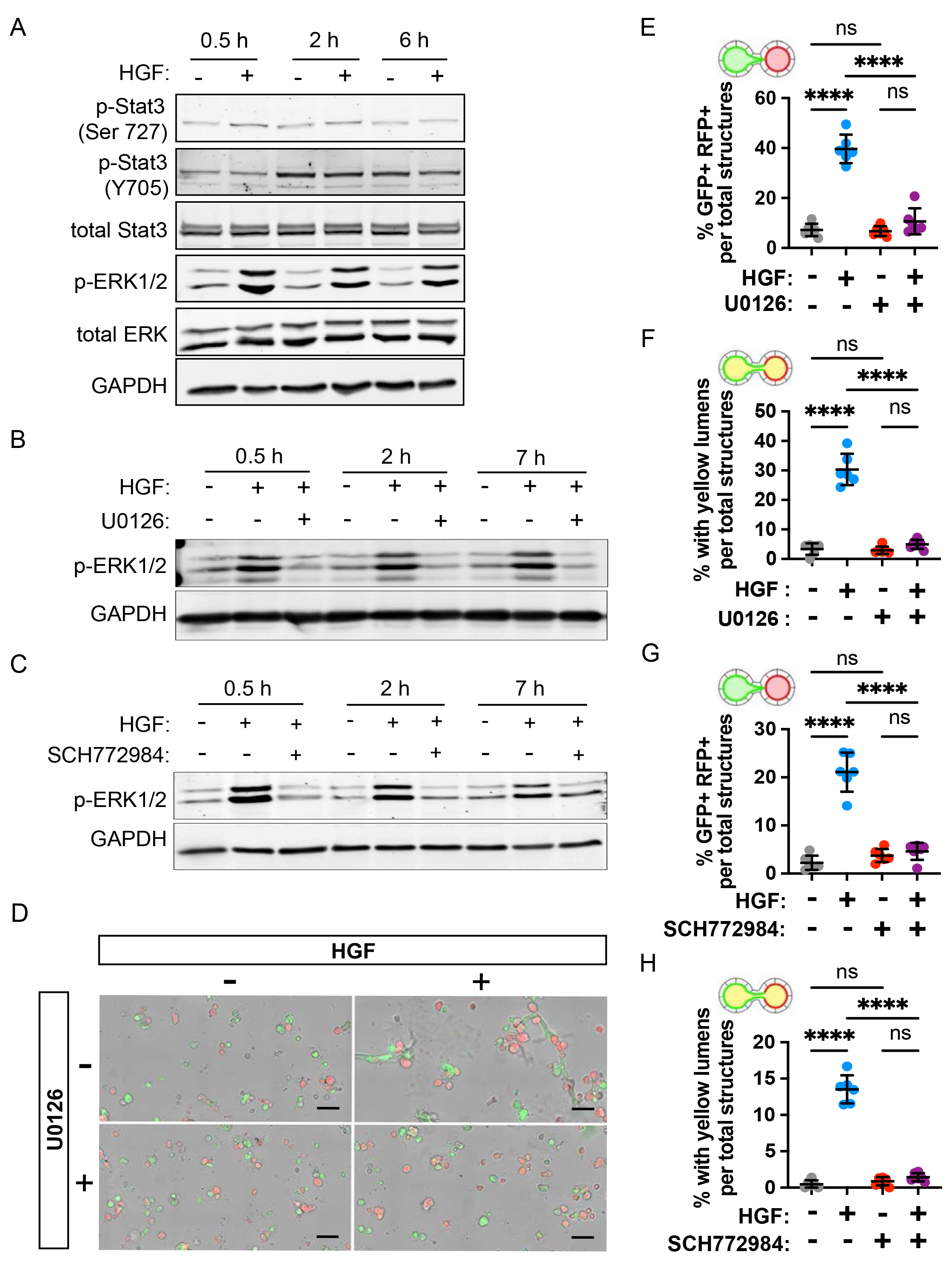
MAPK/ERK signaling is necessary for tubular interconnection. **(A)** Replated MDCK spheroid cells were treated with HGF (10 ng/mL) and lysed after 0.5, 2, and 6 hours. The protein levels of p-Stat3, total Stat3, pERK1/2, and total ERK were measured by western blotting. GAPDH was used as a loading control. **(B,C)** Replated MDCK spheroids were treated with MEK1/2 inhibitor, U0126 (10 µM), and ERK inhibitor, SCH772984 (5 µM), 30 min prior to HGF (10 ng/mL) addition and cells lysed at 0.5, 2, and 7 hours. Protein level of p-ERK1/2 was measured by western blotting, and GAPDH was used as a loading control. **(D-H)** Replated spheroids were treated with either HGF (10ng/mL), U0126 (10 µM), SCH772984 (5 µM) or a combination, for two days. (D) Representative images of replated spheroids after U0126 treatment period. Scale bars = 200 µm. (E, G) Percentage of GFP^+^-RFP^+^ structures induced by HGF was reduced by U0126 and SCH772984. (F, H) Percentage of yellow-lumen structures induced by HGF was reduced by U0126 and SCH772984. Data are representative of at least 2 independent experiments performed in triplicate. Measures of interconnection (E-H) were compared by two-way ANOVA, *P*&lt;.0001****.

To determine if the HGF-induced tubule interconnection was dependent on MEK/ERK activation, we treated the spheroids with an inhibitor of MEK (U0126) or an inhibitor of ERK1/2 (SCH772984). Both inhibitors fully blocked the HGF-induced increase in ERK1/2 phosphorylation (**Fig 5B,C**). To assess the necessity of these pathways in anastomosis, we cultured replated MDCK spheroid cells expressing GFP-Podxl^+^ and RFP-Podxl^+^. The proportion of lumen fusion increased significantly (p < 0.0001) from 3.4% to 30.3% when HGF was added in the culture medium, indicating tubular interconnection between spheroids (**Fig 5D,F**). Spheroids with dual fluorescence and lumen fusion were not observed in the presence of inhibitors of MEK (U0126) or ERK1/2 (SCH772984) (**Fig 5E-H**). Together, these results suggest that stimulation of tubular anastomosis by HGF is mediated by the ERK1/2 signaling pathway.

### Matrix metalloproteases are required for tubular and lumenal fusion

HGF has previously been shown to activate matrix metalloproteases (MMPs) and promote tubule development (54). To test the role of MMPs in mediating HGF-induced tubule anastomosis, we examined the activities of MMP9 and MMP2 using gelatin zymography analysis in our cultured MDCK cells. The amount of active form of extracellular MMP9 increased significantly in response to the HGF treatment over time while MMP2 activity was unchanged (**Fig 6A-D**). These data suggest a potential role of MMP9 in anastomosis. (**Fig 6C,D**). To test this directly, we utilized ilomastat which is known to broadly inhibit MMP function (57, 58). Without ilomastat in the culture medium, HGF promoted tubule and lumen fusion as shown by dual florescence of MDCK structures and lumens (**Fig 6E-G**). Treatment with ilomastat reduced anastomosis at the level of structural interconnection and lumen fusion (**Fig 6E-G**). Collectively, these data confirm roles for MEK-ERK and MMPs downstream of HGF in regulating tubular interconnection.

**Figure 6.**
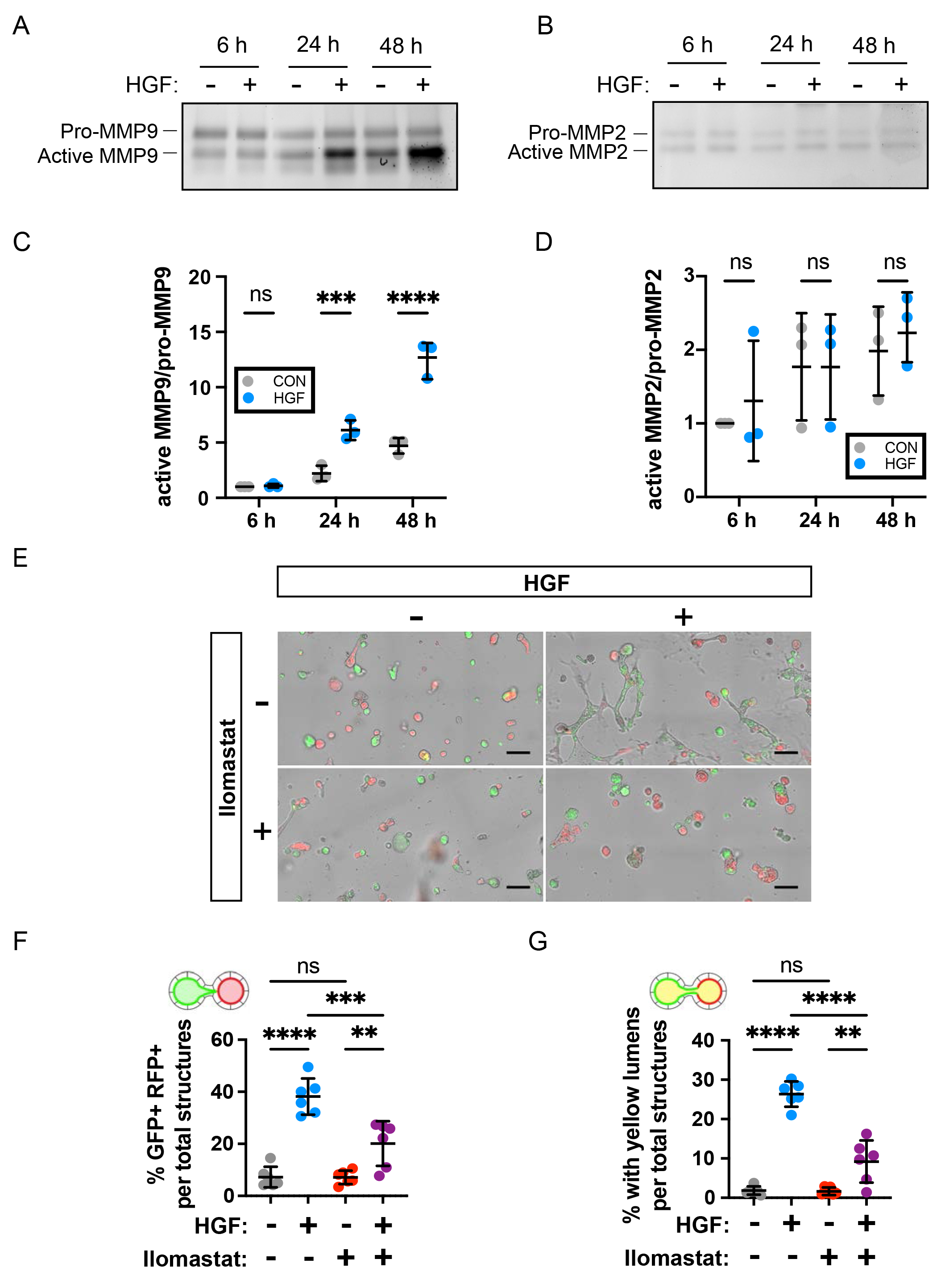
Broad spectrum inhibition of MMPs reduces lumenal interconnection. **(A, B)** Replated 3D MDCK cells were treated with HGF (10 ng/mL) and cultured in serum free medium. Conditioned media was collected at 6, 24, 48 hours and extracellular MMP9 and MMP2 were analyzed by gelatin zymography assay. **(C, D)** Quantification of active MMP9/pro-MMP9 activity and active MMP2/pro-MMP2. Data are expressed in arbitrary units, with the value = 1 corresponding to the activity of 6 h in untreated cells. Measures of quantification were compared by two-way ANOVA, *P*&lt;.0001****, *P*&lt;.001***. **(E-G)** Replated spheroids were treated with HGF (10 ng/mL), ilomastat (GM6001, 50 nM), or a combination, for two days. (E) Representative images of replated spheroids after treatment period. Scale bars = 200 µm. The percentage of GFP^+^-RFP^+^ structures (F) and ‘yellow’ lumen structures (G) from HGF induction was reduced by ilomastat treatment. Data are representative of at least two independent experiments performed in triplicate. Measures of interconnection were compared by two-way ANOVA, *P*&lt;.0001****, *P*&lt;.001***, *P*&lt;.01**.

### HGF promotes tubule interconnection in kidney explants

To investigate whether the mechanisms of HGF-induced anastomosis from an *in vitro* epithelial monoculture system can be extrapolated to a complex multicellular tissue, we examined *ex vivo* culture explants of embryonic day 12.5 (E12.5) kidneys from *Ksp-cre ^Tg/+^;Rosa-tdTomato^f/+^* mice. *Ksp-cre* induces recombination in ureteric bud epithelia (59) and *Rosa td-Tomato* is a reporter allele encoding an RFP variant (60), thereby enabling fluorescent visualization of the cre-recombined ureteric bud. E12.5 kidneys were dissected, and each pair of kidneys was plated with their cortices abutting the other, an orientation that brought their nephrogenic zones and ureteric buds in close proximity. Pairs were cultured with or without HGF (100 μg/mL) for 72 hours, at which time the explants were fixed, cleared, and stained for a marker of tubule lumens, Par6. They were then imaged by confocal microscopy through the full thickness of the explant. Initial experiments did not demonstrate lumenal fusion between ureteric buds (UB), or between ureteric bud (UB) and renal vesicle (RV) of adjacent kidney explants.

We hypothesized that the presence of mesenchymal cells in the cortex of embryonic kidneys created a physical barrier between epithelia that prevented interactions and subsequent anastomosis. To increase the accessibility of tubules between two embryonic kidneys, we treated explants with collagenase A (2.5 mg/mL) for 1, 1.5, and 2 min at 37℃ which removed some of the mesenchymal cells surrounding the kidneys. Embryonic kidneys treated for more than 1.5 minutes did not develop well (not shown) and were not used in subsequent experiments. Using this strategy (**Fig 7A-E**), we found that HGF promotes UB-UB anastomosis between embryonic kidneys treated with collagenase A for 1 min. Staining with lumenal marker Par6 indicated that not only were the tubules interconnected, but their lumens had fused as well (**Fig 7E**). Treatment of explants with HGF or collagenase alone did not induce interconnection (**Fig 7B,C**). Similar interconnections of UB-RV and RV-RV were not observed, which may be due to their positioning and lack of close proximity. Interestingly, where an explant was positioned correctly such that the ureter of one kidney was abutting the cortex of the other, this resulted in anastomosis between the UB and ureter (**Fig S3**). Together these results suggest that extrinsic manipulation of tubules *ex vivo* is sufficient to tubule interconnection and that HGF is a key factor to promote tubule interconnection.

**Figure 7.**
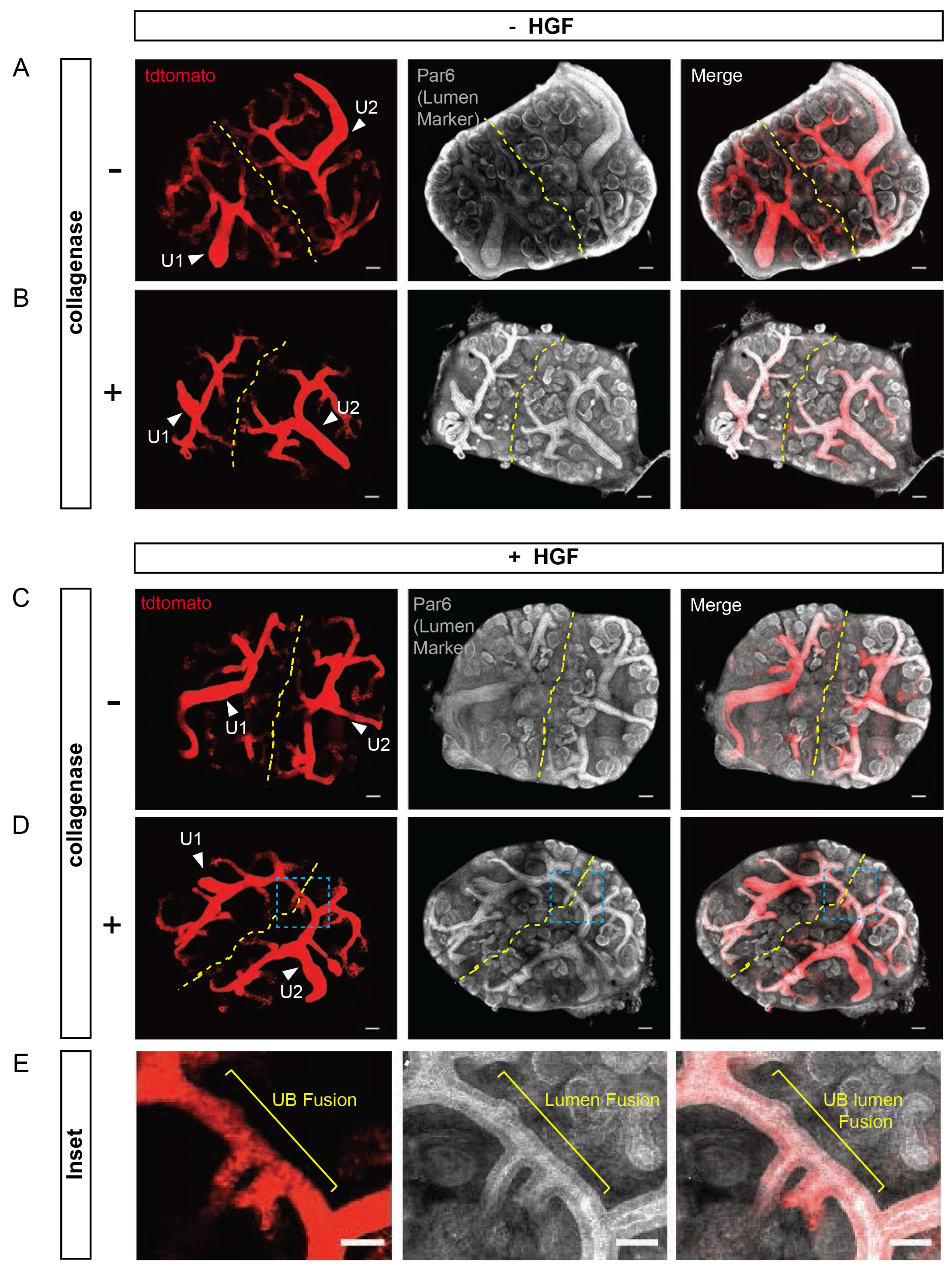
HGF promotes extrinsic UB-UB fusion in embryonic kidney explants. Two E12.5 kidney explants (*Ksp-Cre^Tg/+^;Rosa-tdTomato^f^/^+^*) were co-plated and treated +/- collagenase (2.5 mg/mL) for 1 min at 37℃ and then plated +/- HGF (100 µg/mL) for 72 hours. Immunofluorescence of UB (Red) and ⍺-Par6b (White), which marks the tubule lumens, are shown. **(A, C)** Kidney explants with and without HGF. **(B, D)** Collagenase-treated explants with and without HGF. UB- UB connection is present in D. **(E)** The inset image from D shows the lumen interconnection. Arrowheads indicate ureter (U1, U2). UB: ureteric bud. Scale bars: 100 µm (A-D); 50µm (E). Similar results were obtained in 4 of 5 experiments.

## DISCUSSION

Fusion of epithelial tissues is an essential event in embryonic development and represents a significant barrier in tissue engineering applications. Our current understanding of the process is mainly based on studies from developmental model systems (31, 61–63). During tubule anastomosis in the developing kidney, there is dissolution of intervening extracellular matrix (27, 64), invasion of the ureteric epithelia by the distal renal vesicle (20), reorganization of the cells, formation of new cellular junctions and re-establishment of polarity (21). Despite careful descriptive analyses of these processes, the mechanisms that lead to endogenous tubular anastomosis have remained elusive, due in part to the complexity of the developing organ, which has numerous, interdependent morphogenetic processes. Use of a simple, *in vitro* model offers advantages in this regard. Here we have developed a quantitative *in vitro* assay to visualize and quantify epithelial tubular anastomosis, and successfully used it to identify HGF as a promising candidate that promotes tubule anastomosis and lumen fusion.

To gain insight into how one might promote tubule interconnection ex vivo, we investigated the molecular interactions associated with endogenous tubule anastomosis during kidney development. Using single-cell transcriptomic data from E18.5 mouse kidneys (38, 39), we identified ligand-receptor interactions at sites of tubule anastomoses. Among these interactions, hepatocyte growth factor (HGF) emerged as a highly significant ligand produced by the nephrogenic stroma, with MET and ST14 identified as prominent receptors in the ureteric bud. We chose to focus on HGF due to its solubility and established role as a tubular morphogen. In addition, the HGF receptor MET is widely expressed in adult kidney tubules, making HGF an attractive candidate for additional translational studies. HGF has been shown to exert myriad effects on cell behavior *in vitro* (51, 52, 54, 65). Notably, it promotes the outgrowth of delicate basal cell protrusions from polarized epithelia and induces a partial epithelial-to-mesenchymal transition (66), both critical for anastomosis. Our studies demonstrate that exogenous HGF robustly promotes tubule anastomosis both *in vitro* and *ex vivo* in kidney tubules. Additionally, our data indicate that HGF-induced tubule anastomosis is mediated through MAPK signaling and MMP-dependent pathways.

The role of HGF and its effector signaling pathways (i.e. STAT3, MAPK, MMP activity) have been well studied in models of epithelial tubule formation. The significance of the current work is that it extends knowledge of HGF function to include tubule anastomosis, which has clear translational applications for organoid-based therapies, and it begins to elucidate the processes and signaling pathways required for anastomosis. Not surprisingly, some of these HGF-induced processes are similarly required in both tubule formation and anastomosis, including the essential roles for MAPK signaling. Intriguingly, although HGF/MAPK is known to induce cell proliferation (54), our studies revealed that there is no requirement for cell proliferation in tubule anastomosis. This implies that HGF-induced anastomosis is not secondary to the increased size of HGF-treated spheroids, which would be expected to increase direct contact. It also implies that the transient perturbations of the basement membrane that occur in dividing cells (66) do not contribute.

Previous work has shown that HGF promotes induction of various MMPs in several epithelial cell types (54, 67, 68). Indeed, in our assay, we found that HGF treatment increased MMP-9 activity. To test the necessity for MMP activity in inducing anastomosis, we used the broad-spectrum MMP inhibitor ilomastat. Our results demonstrate that MMP inhibition is sufficient to block tubule anastomosis by HGF, suggesting that collagen degradation and matrix invasion and/or another MMP function, such as re-establishment of epithelial polarization and differentiation (54), is required for tubule and luminal anastomosis.

To validate the biological significance of the *in vitro* results, we tested the ability of HGF to mediate tubule anastomosis in adjacent, explanted embryonic kidneys. Initial experiments were unsuccessful, perhaps due to the lack of close proximity between tubules since several layers of stromal cells intervene between the tubules of adjacent kidneys. A brief treatment with collagenase prior to explant culture served to overcome this hurdle, resulting in robust anastomosis between adjacent ureteric bud tubules. Collagenase may have contributed to anastomosis not only by removing stromal cells, but by enhancing dissolution of the tubule basement membranes, which may have been more resistant to turnover than in the *in vitro* culture model. In any event, future attempts to induce tubule anastomosis via other soluble mediators may benefit from the addition of collagenase.

Exogenous anastomosis of renal vesicles to renal vesicles or other tubules in the adjacent kidney was not observed. This could be attributed to the distance (and intervening cells) between these tubules; however, it may also be explained by the localization of MET to the ureteric bud, and not renal vesicles/nephron. Future experiments to examine these interactions as well as those in other tubule types, while beyond the scope of the current study, will be of great interest. It is important to note that while HGF is known to robustly induce tubulogenesis and anastomosis *in vitro* and ex vivo, inactivation of HGF or its receptor, MET, does not cause observable defects in kidney development, although defects in liver, limb muscles and placental development are present (69–71). Thus, HGF is not required for these processes to occur *in vivo*. This suggests that either HGF can trigger an event that it normally does not affect or, alternatively, that other morphogens with redundant activities are able to compensate for HGF-mediated signaling in epithelia. Support for the later comes from studies in which HGF-blocking antibodies abrogate branching tubulogenesis and nephron formation in kidney explants (72, 73). In addition, EGFR ligands such as TGF-alpha are capable of inducing branching tubulogenesis in MET knockout cells as well as in mIMCD3 cells, where they act in part through MMP activity (74). Consistent with this, mice with deletion of *Met* from the ureteric bud tubules have increased EGFR (75). In further support of functional redundancy, a major downstream signaling cascade of HGF-MET is MAPK, and MAPK signaling can be activated by several ligands in the kidney in addition to HGF. For example, GDNF/RET signaling, which is essential for UB branching and nephron formation, acts in part through MAPK (76). Pharmacologic inhibition of MAPK signaling in kidney explants leads to abrogated ureteric bud branching (77), and genetic ablation of MAPK signaling in the ureteric bud via dual deletion of MEK1/2 prevents ureteric bud branching, causing severe kidney hypodysplasia (78). Collectively, these results suggest an important role for HGF/MET and MAPK signaling in tubule formation/branching, one whose role is masked by functional redundancy by other ligands and/or signaling pathways. Furthermore, although not tested here, one can imagine that soluble ligands other than HGF might similarly affect tubule anastomosis.

In conclusion, organoid biology is becoming increasingly relevant as the promise of autologous, lab-grown, transplantable tissues gives new hopeful directions to transplant medicine (6, 79–81). An essential part of that endeavor is ensuring that interconnections occur between transplanted and host tissues, something that has yet to be achieved. Our cell-based assay and bioinformatic approach can be used to generate novel hypotheses and test additional promising candidates for their effects on tubule anastomosis in a quantitative and relatively efficient manner. Together they will serve as a useful resource for additional studies on tubule interconnection.

## ACKNOWLEDGEMENTS

We would like to thank Drs. V. Patel (UTSW), H. Zhu (UTSW), and D. Bryant (U. Glasgow) for helpful comments and feedback. We thank M. Martinez and B. Felan for assistance with preliminary experiments. This work was supported by NIH RC2DK125960, UC2DK126021, P30DK079328, and the Carolyn R. Bacon Distinguished Professorship in Medical Science and Education (DKM).

## AUTHOR CONTRIBUTIONS

Conceptualization: DKM; Methodology: DKM, ILG, SO; Software: CC, TC; Validation: SO; Supervision: DKM; Data analysis: ILG, SO; Investigation: SO, ILG, JT; Writing of the first draft: DKM, ILG, SO. All authors reviewed and revised the manuscript.

## METHODS

### Cells lines and spheroid culture

MDCKII cells were cultured in MEM/EBSS with 5% FBS, 1% P/S, and 15 mM HEPES, and tested for mycoplasma contamination. To generate stable cell lines producing GFP-podocalyxin or RFP- podocalyxin, we cloned canine *Podxl1* into the pQCXIZ retroviral plasmid with a GFP or RFP fused to the N-terminus (a gift from Eric Campeau, Addgene plasmid #22801). We packaged retroviruses containing each vector in HEK293GPG using Mirus Trans-ItLT1 transfection reagent. Viral supernatant was collected for 3 days and then used for cell transduction. Transduced cells were selected with Zeocin (100 µg/ml, Invitrogen, #R25001) for 14 days and then FAC-sorted to enrich fluorescence (top 30%) for further experiments.

### Spheroid Anastomosis Assay

MDCKII spheroids were grown in Matrigel (Corning, #354234) as described previously (82). Briefly, single cell suspensions (4.4x10^4^ cells/mL) of MDCKII GFP-podocalyxin or RFP- podocalyxin were prepared in media with 2% Matrigel and plated on polymerized Matrigel (5 µl/cm^2^). Seventy-two-hour cultures were rinsed with ice-cold PBS+ (PBS, 1mM CaCl2, 0.5 mM MgCl2), and treated with Cell Recovery solution (Corning, #354253) containing 10 U/mL DNase (Promega, #M6101) for 1 hour at 4°C with agitation. Depolymerized slurries were triturated 10 times to release spheroids, mixed 1:1 the GFP-podocalyxin and RFP-podocalyxin spheroids, and centrifuged at 900*g* for 90 seconds at 4°C. Supernatants containing depolymerized Matrigel were discarded. Spheroids were gently resuspended in media with 2% Matrigel and plated in 8-well chamber slides, approximately 4- to 10-fold denser per unit area than the individual seeded cultures. HGF (R&D, #2207), mitomycin-C (Sigma, #M5353), U0126 (Promega, #V112A), SCH772984 (Sellekchem, #S7101), or ilomastat (Selleckchem, #S7157) were added as indicated in experiments. All conditions were performed in triplicate.

### Image acquisition

Baseline spheroid count was determined by tile-scanning replated spheroids in triplicate with Nikon Eclipse Ti widefield microscope three hours after harvest. Spheroids were re-imaged at 24 and 48 hours. ImageJ was used for image analysis. Briefly, calibrated brightfield images were subjected to 100 µm Gaussian blur, adjusted for brightness, thresholded, and converted to binary. Particles identified in binary images were closed, filled, and counted. In a blinded manner, particles were superimposed on fluorescent channels to score GFP^+^-RFP^+^ (dual-positive) structures and lumen spectral overlap (yellow-lumens). Dual-positive, yellow-lumen, and endpoint structure count over baseline were reported as percentages of total structures. Shape determinants such as cross-sectional area was measured and reported as frequency distributions for each treatment condition. Images in figures were linearly adjusted for brightness. All conditions were performed in triplicate and analysis of images was performed in a blinded fashion.

### Immunoblot assay

MDCK spheroids replated onto 12-well plates were treated with 10 µM U0126 or 5 µM SCH772984 for 30 min prior to addition of HGF (10 ng/ml). After the spheroid harvest, total protein was extracted in RIPA buffer (100 mM NaCl, 20 mM NaPO4, pH 7.2, 0.1% SDS, 0.5% sodium deoxycholate, 1% NP40, 50 mM NaF, 4 mM sodium vanadate, protease inhibitor, 1 mM PMSF). The protein was quantified by BCA assay and equivalent amounts of protein were electrophoresed in SDS-PAGE and transferred to nitrocellulose membrane. The membrane was blocked in 5% BSA and probed with respective primary antibodies: p-ERK (Cell Signaling, #9101S), total ERK (Cell Signaling, #4695), p-Stat3(Ser 727, Cell Signaling, #9134), p-Stat3 (Y705, Cell Signaling, #9145s), total Stat3 (Cell Signaling, #9132) and GAPDH. IRDye680RD Donkey anti-rabbit and IRDye800CW Donkey anti-mouse (Licor) were used, and images were scanned using the ChemiDoc MP TM system (Bio-Rad, Hercules, CA, USA).

### Gelatin Zymography

Three-day cultured spheroids were replated onto 12-well plates coated with Matrigel and treated with HGF (10 ng/ml) in serum-free medium. The conditioned media were collected at 2, 24, and 48 hours and gelatin zymography was performed using 7.5% acrylamide gel containing 4 mg/mL gelatin (SIGMA). Briefly, serum-free conditioned media was mixed with 5X non-reducing sample buffer, incubated at 37°C for 20 min, and then subjected to 7.5% SDS-PAGE. The gel was washed for 30 minutes 4 times using renaturing buffer (2.5% Triton X-100, 50 mM Tris, 5 mM CaCl2, 1 µM ZnCl2) and subsequently incubated in developing buffer (1% Triton X-100, 50 mM Tris, 5 mM CaCl2, 1 µM ZnCl2) at 37°C for 2 days. The gels were stained using 0.5% Coomassie Blue solution, then destained in 40% methanol,10% acetic acid. The gels were scanned by ChemiDoc MP TM system (Bio-Rad, Hercules, CA, USA).

### Bioinformatic data, candidate prioritization, and ligand-receptor analyses

We used single cell RNA sequencing datasets of mouse embryonic kidney (E18.5) available on the Gene Expression Omnibus (GEO): <u>GSE155794</u> (35) <u>GSE108291</u> (34). Each batch was processed independently using the scran Bioconductor package (83). Unfiltered feature-barcode matrices were generated by running the CellRanger count pipeline. Cells were called from empty droplets by testing for deviation of the expression profile for each cell from the ambient RNA pool (84). Cells with large mitochondrial proportions, i.e., more than 3 mean-absolute deviations away from the median, were removed. Cells were pre-clustered; a deconvolution method was applied to compute size factors for all cells (85) and normalized log-expression values were calculated. Variance was partitioned into technical and biological components by assuming technical noise was Poisson-distributed and attributing any estimated variance more than that accounted for by a fitted Poisson trend to biological variation. The dimensionality of the data set was reduced by performing principal component analysis and discarding the later principal components for which the variance explained was less than variance attributable to technical noise.

Masking of biological effects by expression changes due to cell cycle phase were mitigated by blocking on this covariate. The cell cycle phase was inferred using the pair-based classifier implemented in the cyclone function of scran. Corrected log-normalized expression counts were obtained by calling the removeBatchEffect from the limma (25605792) Bioconductor package with a design formula including G1 and G2M cell cycle phase scores as covariates. A single set of features for batch correction was obtained by computing the average biological component of variation across batches and retaining those genes with a positive biological component. The batches were rescaled, and log-normalized expression values recomputed after the size factors were adjusted for systemic differences in sequencing depth between batches. Batch effects were corrected by matching mutual nearest neighbors in the high-dimensional expression space (86). The resulting reduced-dimensional representation of the data was used for all subsequent embeddings such as UMAP. Cells were clustered by building a shared nearest neighbor graph (87) and executing the Walktrap algorithm (https://dx.doi.org/10.7155/jgaa.00124). Cluster- specific gene expression was identified with the EIGEN algorithm (36). Putative cell type identities were assigned based on the cluster-specific expression of established marker genes.

Potential ligand-receptor relationships evaluated were those identified by the FANTOM5 project (88). The curated human gene annotations were mapped to mouse orthologues using the AnnotationHub Bioconductor package (https://doi.org/10.18129/B9.bioc.AnnotationHub). For each pair of cell types postulated to serve as source or target, respectively, for a ligand-receptor interaction, the expression of ligands in the former and receptors in the latter were ranked using EIGEN. The ligand-receptor interactions were then ranked by the mean of the ligand and receptor rankings. For a given pair of interacting cell types, ligands are ranked in descending order by expression specificity in the source population on the left-hand side and receptors ranked in descending order of expression specificity in the target population on the right-hand side. A line segment connects the ligands to receptors of interacting pairs. The higher the “rung of the ladder”, the more specific the ligand-receptor interaction.

### Statistics

Measures of anastomosis (dual-positive and ‘yellow’ lumen structures, cross-sectional area, and structure number) were compared by Student’s t-test or two-way ANOVA as indicated in each experiment.

## SUPPLEMENTAL INFORMATION TITLES AND LEGENDS

**Supplemental Table S1. Ligand-receptor analysis in E18.5 embryonic mouse kidneys.** See methods for experimental details. Top 100 ligand-receptor pairs for each combination of interacting cell types are listed. Full interaction results are available upon request.

Supplemental Figure S1. **HGF-induced anastomosis is dose dependent.** Three-day spheroids were replated and treated with HGF (0 - 30 ng/mL). Structures were imaged and counted after two days of treatment. Data are representative of at least five independent experiments performed in triplicate.

Supplemental Figure S2. **Mitomycin C inhibits proliferation and maintains spheroid size with minimal cell death. (A)** Three-day old MDCK spheroids were treated with Mitomycin C for two days. Representative brightfield images are shown. **(B)** Immunofluorescence of proliferating (EdU-AlexaFluor647, magenta) and dying cells (activated Caspase 3, green). Nuclei are counter- stained with Hoechst 33342. **(C)** Quantification of cross-sectional area of spheroids in (A). Dashed line and grey regions indicate average ± 1 and 2 SD of three-day old spheroids. **(D)** Cross- sectional area as in (C) with narrowed treatment range.

Supplemental Figure S3. **HGF promotes extrinsic UB-ureter fusion in kidney explants.** Two kidney explants (*Ksp-Cre^Tg/+^;Rosa-tdTomato^f/+^*and *Ksp-Cre^Tg/+^*) were co-plated and treated +/- collagenase (2.5 mg/mL) for 1 min at 37℃ and then plated +/- HGF (100 µg/mL) for 72 hours. Immunofluorescence of UB (Red) and ⍺-Par6b (White), which marks the tubule lumens, is shown. **(A, C)** Kidney explants with and without HGF. **(B, D)** Collagenase-treated explants with and without HGF. UB-ureter connection is present in D only. **(E)** Inset image from D shows lumen interconnection. Arrowheads indicate ureters (U1, U2). Scale bars: 100 µm (A-D); 50 µm (E).

