## Supplementary figures and images for "Epithelial tubule interconnection driven by HGF-Met signaling in the kidney"

### Supplemental Figures S1-2

Supplemental Figure S1.

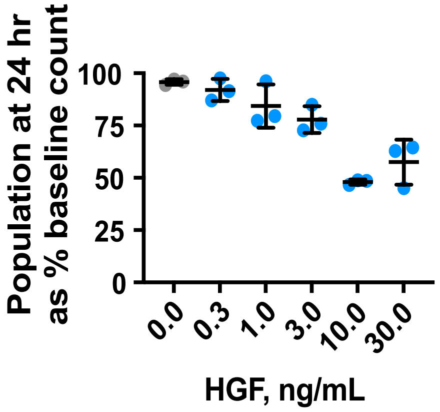

Supplemental Figure S2.

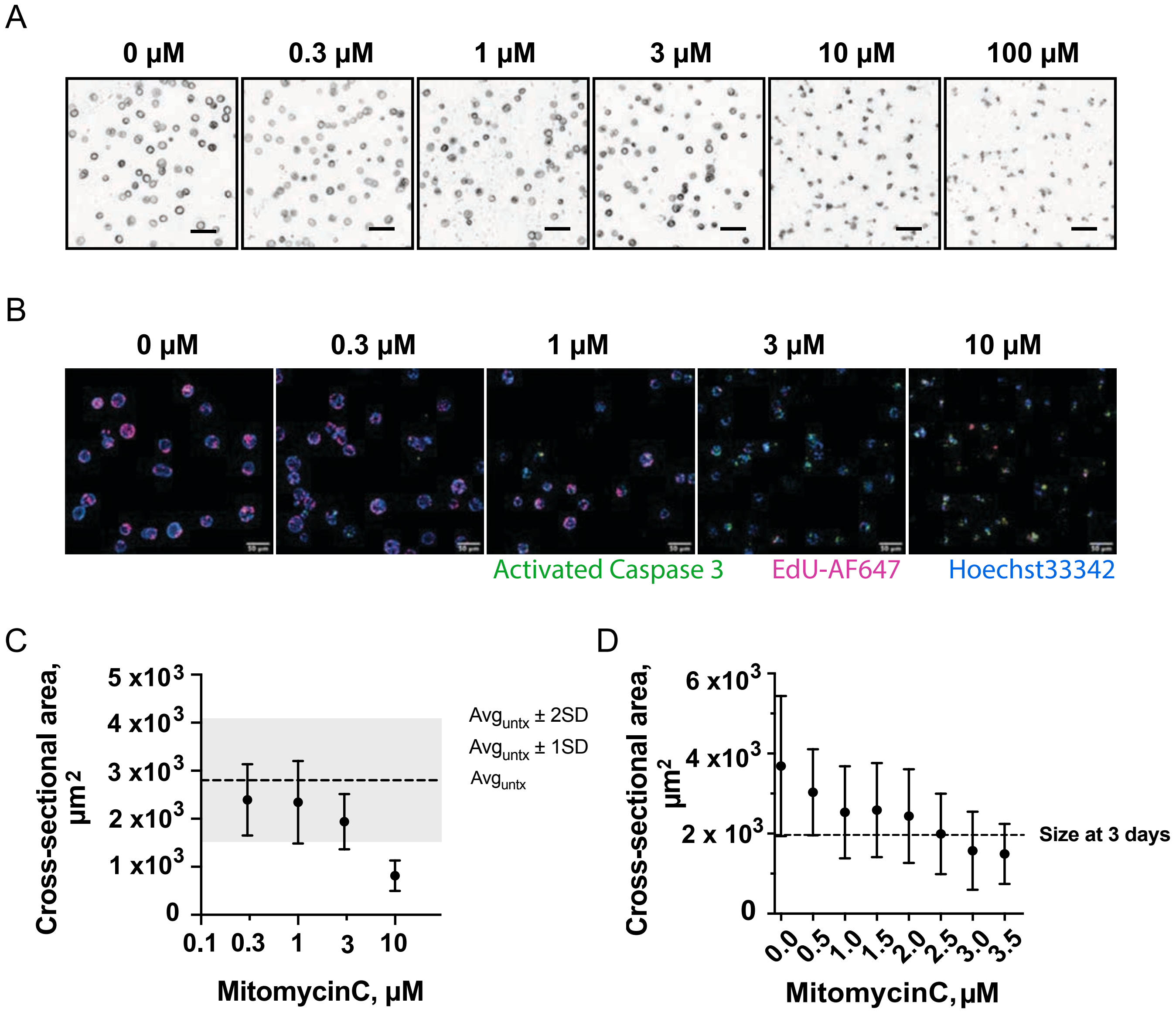

Supplemental Figure S3 .

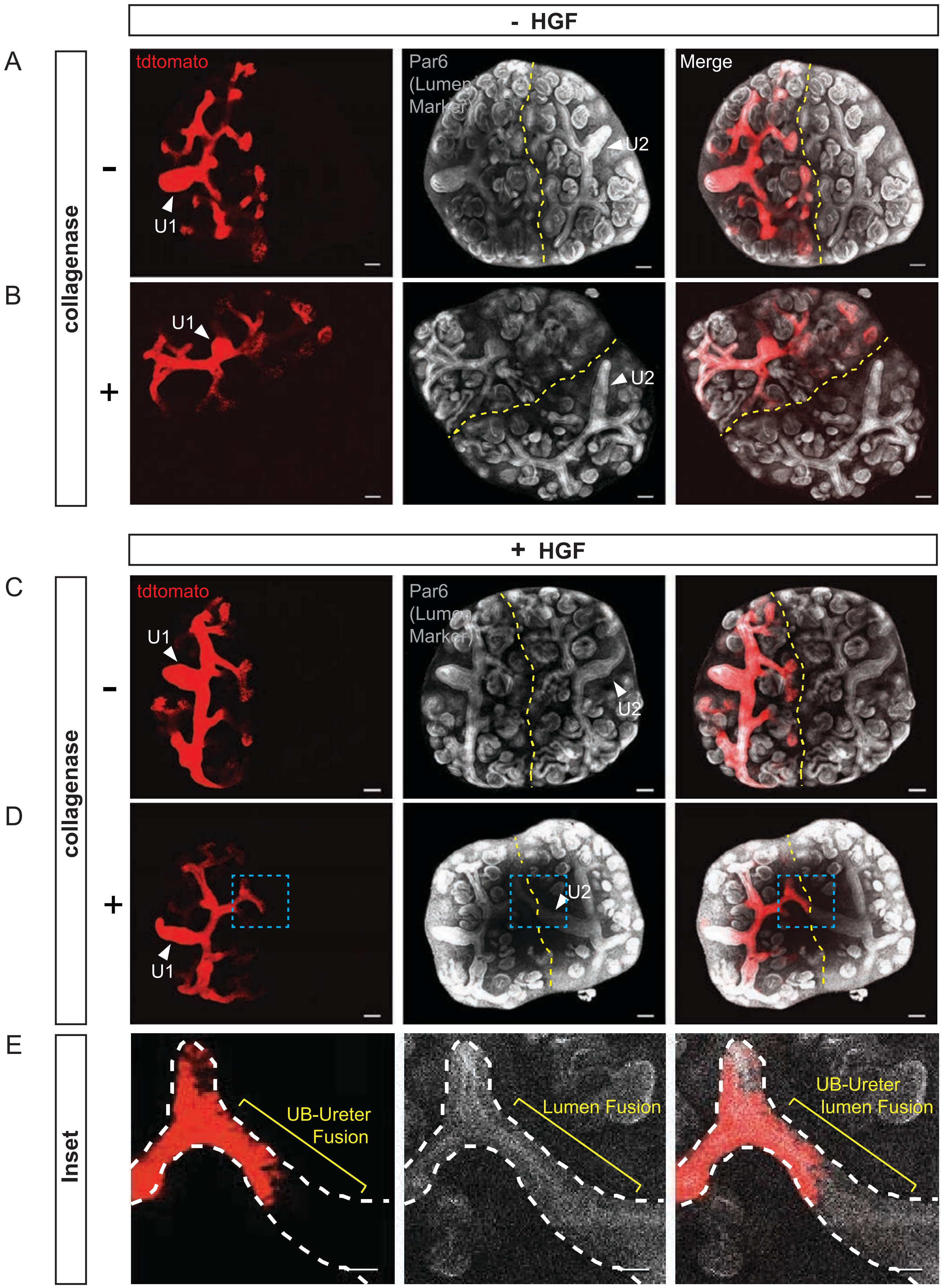
