## Supplemental Table S1 for "Epithelial tubule interconnection driven by HGF-Met signaling in the kidney"

Supplemental Table 1 SP1-RV

| Ligand | Receptor | Ligand rank | Receptor rank | Interaction rank |
|---|---|---|---|---|
| **Mfap2** | Notch1 | 56 | 157 | 1 |
| **Dlk1** | Notch1 | 64 | 157 | 2 |
| **Mfap5** | Notch1 | 66 | 157 | 3 |
| **Fgf2** | Fgfr1 | 409 | 26 | 4 |
| **Ncam1** | Fgfr1 | 531 | 26 | 5 |
| **Efna5** | Epha4 | 533 | 113 | 6 |
| **Ntf5** | Ngfr | 534 | 175 | 7 |
| **Fgf2** | Sdc1 | 409 | 329 | 8 |
| **App** | Ngfr | 650 | 175 | 9 |
| **Lpl** | Sdc1 | 522 | 329 | 10 |
| **Nid1** | Col13a1 | 712 | 158 | 11 |
| **Cthrc1** | Fzd3 | 523 | 380 | 12 |
| **Fgf2** | Fgfrl1 | 409 | 495 | 13 |
| **Adam17** | Notch1 | 790 | 157 | 14 |
| **Inha** | Acvr2b | 713 | 281 | 15 |
| **App** | Gpc1 | 650 | 712 | 16 |
| **Ntn1** | Unc5c | 1 | 1383 | 17 |
| **Fgf2** | Fgfr2 | 409 | 1003 | 18 |
| **Lama4** | Itga6 | 26 | 1394 | 19 |
| **Ntn1** | Unc5b | 1 | 1437 | 20 |
| **Efnb1** | Epha4 | 1359 | 113 | 21 |
| **Ncam1** | Fgfr2 | 531 | 1003 | 22 |
| **Efna5** | Epha7 | 533 | 1044 | 23 |
| **Dlk1** | Notch2 | 64 | 1618 | 24 |
| **Rspo3** | Lgr4 | 3 | 1708 | 25 |
| **Ecm1** | Cachd1 | 140 | 1599 | 26 |
| **Lamc1** | Itga6 | 349 | 1394 | 27 |
| **Psap** | Celsr1 | 1311 | 472 | 28 |
| **Col6a1** | Itga6 | 443 | 1394 | 29 |
| **Efna5** | Ephb1 | 533 | 1571 | 30 |
| **Fgf9** | Fgfr1 | 2224 | 26 | 31 |
| **Lrpap1** | Sorl1 | 390 | 1996 | 32 |
| **Lama4** | Itga3 | 26 | 2445 | 33 |
| **Wnt5a** | Fzd4 | 758 | 1769 | 34 |
| **Rspo3** | Lgr6 | 3 | 2551 | 35 |
| **Col5a2** | Ddr1 | 138 | 2425 | 36 |
| **Nid2** | Col13a1 | 2419 | 158 | 37 |
| **Col1a1** | Ddr1 | 210 | 2425 | 38 |
| **Tgfb3** | Tgfbr1 | 17 | 2619 | 39 |
| **Reln** | Itga3 | 196 | 2445 | 40 |
| **Wnt5a** | Ryk | 758 | 1954 | 41 |
| **Ncam1** | Robo1 | 531 | 2214 | 42 |
| **Fn1** | Col13a1 | 2611 | 158 | 43 |
| **Lamc1** | Itga3 | 349 | 2445 | 44 |
| **Efnb1** | Ephb1 | 1359 | 1571 | 45 |
| **Igf2** | Igf1r | 199 | 2769 | 46 |
| **Igf1** | Igf1r | 358 | 2769 | 47 |
| **Nid1** | Itga3 | 712 | 2445 | 48 |
| **Thbs1** | Sdc1 | 2834 | 329 | 49 |
| **Col11a1** | Ddr1 | 742 | 2425 | 50 |
| **Rspo3** | Sdc4 | 3 | 3185 | 51 |
| **Hgf** | Sdc1 | 2892 | 329 | 52 |
| **Fgf18** | Fgfr1 | 3198 | 26 | 53 |
| **Apoe** | Sorl1 | 1228 | 1996 | 54 |
| **Fgf9** | Fgfr2 | 2224 | 1003 | 55 |
| **App** | Slc45a3 | 650 | 2593 | 56 |
| **Tfpi** | Sdc4 | 62 | 3185 | 57 |
| **Ntn1** | Neo1 | 1 | 3329 | 58 |
| **Lrpap1** | Ldlr | 390 | 2949 | 59 |
| **Col3a1** | Ddr1 | 990 | 2425 | 60 |
| **Gm11808** | Notch1 | 3293 | 157 | 61 |
| **Dlk1** | Notch4 | 64 | 3487 | 62 |
| **Fgf2** | Sdc4 | 409 | 3185 | 63 |
| **Tgfb2** | Tgfbr1 | 996 | 2619 | 64 |
| **Lrpap1** | Lrp2 | 390 | 3305 | 65 |
| **Rarres2** | Gpr1 | 444 | 3262 | 66 |
| **B2m** | Tfrc | 2338 | 1399 | 67 |
| **Lama5** | Sdc1 | 3446 | 329 | 68 |
| **Lamb1** | Itga6 | 2411 | 1394 | 69 |
| **Apoe** | Scarb1 | 1228 | 2581 | 70 |
| **Lpl** | Lrp2 | 522 | 3305 | 71 |
| **Gdnf** | Gfra1 | 343 | 3510 | 72 |
| **Tgm2** | Sdc4 | 670 | 3185 | 73 |
| **Timp2** | Itga3 | 1435 | 2445 | 74 |
| **Thbs2** | Itga6 | 2491 | 1394 | 75 |
| **Rgma** | Neo1 | 560 | 3329 | 76 |
| **Hspg2** | Col13a1 | 3760 | 158 | 77 |
| **Il2** | Ngfr | 3746 | 175 | 78 |
| **Myoc** | Fzd3 | 3613 | 380 | 79 |
| **Fn1** | Itga6 | 2611 | 1394 | 80 |
| **Ncam1** | Gfra1 | 531 | 3510 | 81 |
| **Lamc1** | Itgb4 | 349 | 3720 | 82 |
| **Hspg2** | Sdc1 | 3760 | 329 | 83 |
| **Myoc** | Fzd10 | 3613 | 486 | 84 |
| **Sfrp1** | Fzd6 | 36 | 4071 | 85 |
| **Efna4** | Epha4 | 3996 | 113 | 86 |
| **Wnt5a** | Fzd7 | 758 | 3385 | 87 |
| **Nid1** | Ptprf | 712 | 3463 | 88 |
| **Apoe** | Ldlr | 1228 | 2949 | 89 |
| **Fgf18** | Fgfr2 | 3198 | 1003 | 90 |
| **Thbs1** | Itga6 | 2834 | 1394 | 91 |
| **Dll4** | Notch1 | 4152 | 157 | 92 |
| **Rspo3** | Fzd8 | 3 | 4361 | 93 |
| **Tfpi** | F3 | 62 | 4372 | 94 |
| **Apoe** | Lrp2 | 1228 | 3305 | 95 |
| **Cthrc1** | Fzd6 | 523 | 4071 | 96 |
| **Rtn4** | Ngfr | 4503 | 175 | 97 |
| **Nrtn** | Gfra1 | 1241 | 3510 | 98 |
| **Wnt5a** | Fzd6 | 758 | 4071 | 99 |
| **Efna3** | Epha4 | 4717 | 113 | 100 |

Supplemental Table 1 SP1-UB

| Ligand | Receptor | Ligand rank | Receptor rank | Interaction rank |
|---|---|---|---|---|
| **Tfpi** | Vldlr | 62 | 29 | 1 |
| **Ntn1** | Neo1 | 1 | 215 | 2 |
| **Reln** | Vldlr | 196 | 29 | 3 |
| **Rspo3** | Lgr5 | 3 | 298 | 4 |
| **Gdnf** | Gfra1 | 343 | 2 | 5 |
| **Gdnf** | Ret | 343 | 3 | 6 |
| **Lrpap1** | Vldlr | 390 | 29 | 7 |
| **Ncam1** | Gfra1 | 531 | 2 | 8 |
| **Fgf2** | Fgfr2 | 409 | 137 | 9 |
| **Lpl** | Vldlr | 522 | 29 | 10 |
| **Ncam1** | Fgfr2 | 531 | 137 | 11 |
| **Rgma** | Bmpr1b | 560 | 130 | 12 |
| **Rgma** | Neo1 | 560 | 215 | 13 |
| **Nid1** | Ptprf | 712 | 352 | 14 |
| **Ntn1** | Unc5b | 1 | 1104 | 15 |
| **Nrtn** | Gfra1 | 1241 | 2 | 16 |
| **Nrtn** | Ret | 1241 | 3 | 17 |
| **Apoe** | Vldlr | 1228 | 29 | 18 |
| **Rspo3** | Sdc4 | 3 | 1432 | 19 |
| **Igf2** | Igf1r | 199 | 1258 | 20 |
| **Dcn** | Met | 446 | 1018 | 21 |
| **Tfpi** | Sdc4 | 62 | 1432 | 22 |
| **Igf1** | Igf1r | 358 | 1258 | 23 |
| **Lrpap1** | Sort1 | 390 | 1356 | 24 |
| **Efna5** | Ephb2 | 533 | 1248 | 25 |
| **Fgf2** | Sdc4 | 409 | 1432 | 26 |
| **Efna5** | Epha4 | 533 | 1380 | 27 |
| **Wnt5a** | Fzd4 | 758 | 1222 | 28 |
| **Lama4** | Itga6 | 26 | 1961 | 29 |
| **Tgm2** | Sdc4 | 670 | 1432 | 30 |
| **Tbc1d24** | Neo1 | 1989 | 215 | 31 |
| **Ltbp3** | Itgb5 | 489 | 1810 | 32 |
| **Lamc1** | Itga6 | 349 | 1961 | 33 |
| **Fgf9** | Fgfr2 | 2224 | 137 | 34 |
| **Efnb1** | Erbb2 | 1359 | 1009 | 35 |
| **App** | Slc45a3 | 650 | 1721 | 36 |
| **Col6a1** | Itga6 | 443 | 1961 | 37 |
| **Cthrc1** | Fzd3 | 523 | 1947 | 38 |
| **Tfpi** | F3 | 62 | 2452 | 39 |
| **Vegfb** | Ret | 2511 | 3 | 40 |
| **Btc** | Erbb2 | 1555 | 1009 | 41 |
| **Efnb1** | Ephb2 | 1359 | 1248 | 42 |
| **Psap** | Sort1 | 1311 | 1356 | 43 |
| **Cntn4** | Ptprg | 377 | 2348 | 44 |
| **Efnb1** | Epha4 | 1359 | 1380 | 45 |
| **Efna5** | Epha1 | 533 | 2389 | 46 |
| **Mfap2** | Notch1 | 56 | 2893 | 47 |
| **Dlk1** | Notch1 | 64 | 2893 | 48 |
| **Mfap5** | Notch1 | 66 | 2893 | 49 |
| **Fgf2** | Fgfrl1 | 409 | 2614 | 50 |
| **Hgf** | St14 | 2892 | 255 | 51 |
| **Apoe** | Scarb1 | 1228 | 2071 | 52 |
| **Rspo3** | Lrp6 | 3 | 3312 | 53 |
| **Fgf18** | Fgfr2 | 3198 | 137 | 54 |
| **Btc** | Erbb3 | 1555 | 1843 | 55 |
| **Gm11808** | Bmpr1b | 3293 | 130 | 56 |
| **Clcf1** | Crlf1 | 3589 | 10 | 57 |
| **Adam17** | Notch1 | 790 | 2893 | 58 |
| **Wnt5a** | Ror1 | 758 | 3115 | 59 |
| **Hgf** | Met | 2892 | 1018 | 60 |
| **Lama4** | Itga3 | 26 | 3958 | 61 |
| **Reln** | Itga3 | 196 | 3958 | 62 |
| **Inha** | Acvr2a | 713 | 3452 | 63 |
| **Lama5** | Bcam | 3446 | 772 | 64 |
| **Thbs1** | Sdc4 | 2834 | 1432 | 65 |
| **Lamc1** | Itgb4 | 349 | 3929 | 66 |
| **Gm11808** | Erbb2 | 3293 | 1009 | 67 |
| **Lamc1** | Itga3 | 349 | 3958 | 68 |
| **Lamb1** | Itga6 | 2411 | 1961 | 69 |
| **Gnai2** | Adora1 | 4185 | 214 | 70 |
| **Dcn** | Erbb4 | 446 | 3994 | 71 |
| **Dkk2** | Lrp6 | 1132 | 3312 | 72 |
| **Thbs2** | Itga6 | 2491 | 1961 | 73 |
| **Rspo1** | Lgr5 | 4273 | 298 | 74 |
| **Fn1** | Itga6 | 2611 | 1961 | 75 |
| **Ltbp1** | Itgb5 | 2782 | 1810 | 76 |
| **Ntn1** | Unc5c | 1 | 4592 | 77 |
| **App** | Gpc1 | 650 | 3959 | 78 |
| **Nid1** | Itga3 | 712 | 3958 | 79 |
| **Wnt5a** | Ryk | 758 | 3918 | 80 |
| **Wnt5a** | Lrp5 | 758 | 3923 | 81 |
| **Lrpap1** | Sorl1 | 390 | 4292 | 82 |
| **Igfbp4** | Lrp6 | 1415 | 3312 | 83 |
| **Sfrp1** | Fzd6 | 36 | 4716 | 84 |
| **Adam17** | Erbb4 | 790 | 3994 | 85 |
| **Thbs1** | Itga6 | 2834 | 1961 | 86 |
| **Fgf2** | Sdc1 | 409 | 4403 | 87 |
| **Myoc** | Fzd4 | 3613 | 1222 | 88 |
| **Clec11a** | Kit | 153 | 4751 | 89 |
| **Thbs1** | Scarb1 | 2834 | 2071 | 90 |
| **Lpl** | Sdc1 | 522 | 4403 | 91 |
| **Tgfb3** | Tgfbr1 | 17 | 4970 | 92 |
| **Apoe** | Lrp5 | 1228 | 3923 | 93 |
| **Efna5** | Ephb6 | 533 | 4629 | 94 |
| **Fbn1** | Itgb6 | 858 | 4308 | 95 |
| **Cthrc1** | Fzd6 | 523 | 4716 | 96 |
| **Ndp** | Fzd4 | 4032 | 1222 | 97 |
| **Gnai2** | Unc5b | 4185 | 1104 | 98 |
| **Fgf7** | Fgfr2 | 5191 | 137 | 99 |
| **Efna4** | Epha4 | 3996 | 1380 | 100 |

Supplemental Table 1 SP2-RV

| Ligand | Receptor | Ligand rank | Receptor rank | Interaction rank |
|---|---|---|---|---|
| **Gdnf** | Gfra1 | 11 | 42 | 1 |
| **Rspo3** | Lgr5 | 493 | 87 | 2 |
| **Dkk1** | Lrp6 | 1 | 670 | 3 |
| **Dkk2** | Lrp6 | 143 | 670 | 4 |
| **Reln** | Vldlr | 35 | 941 | 5 |
| **Rspo3** | Lrp6 | 493 | 670 | 6 |
| **Rspo3** | Fzd8 | 493 | 778 | 7 |
| **Fgf2** | Sdc1 | 353 | 953 | 8 |
| **Dlk1** | Notch1 | 149 | 1182 | 9 |
| **Fgf2** | Fgfr4 | 353 | 1016 | 10 |
| **Fgf2** | Fgfr1 | 353 | 1211 | 11 |
| **Mfap2** | Notch1 | 411 | 1182 | 12 |
| **Efna2** | Epha4 | 1615 | 15 | 13 |
| **Gdf11** | Acvr2b | 752 | 887 | 14 |
| **Sema3a** | Plxna4 | 217 | 1457 | 15 |
| **Fgf2** | Gpc4 | 353 | 1359 | 16 |
| **Efna3** | Epha4 | 1710 | 15 | 17 |
| **Nid1** | Ptprf | 1216 | 568 | 18 |
| **Lama4** | Itga6 | 1599 | 188 | 19 |
| **Rspo3** | Lgr4 | 493 | 1322 | 20 |
| **Slit2** | Robo1 | 1772 | 56 | 21 |
| **Fn1** | Itga6 | 1677 | 188 | 22 |
| **Cxcl12** | Sdc4 | 90 | 1887 | 23 |
| **Efna2** | Epha7 | 1615 | 409 | 24 |
| **Efna3** | Epha7 | 1710 | 409 | 25 |
| **Gm11808** | Notch1 | 965 | 1182 | 26 |
| **Efna4** | Epha4 | 2184 | 15 | 27 |
| **Fgf2** | Sdc4 | 353 | 1887 | 28 |
| **Igfbp4** | Lrp6 | 1633 | 670 | 29 |
| **Col6a1** | Itga6 | 2165 | 188 | 30 |
| **Rspo3** | Sdc4 | 493 | 1887 | 31 |
| **Cxcl12** | Cxcr4 | 90 | 2292 | 32 |
| **Igfbp4** | Fzd8 | 1633 | 778 | 33 |
| **Efna4** | Epha7 | 2184 | 409 | 34 |
| **Hgf** | St14 | 1795 | 883 | 35 |
| **Slit2** | Sdc1 | 1772 | 953 | 36 |
| **Fgf2** | Fgfr2 | 353 | 2386 | 37 |
| **Hgf** | Sdc1 | 1795 | 953 | 38 |
| **Gnai2** | Adra2a | 743 | 2153 | 39 |
| **Efnb3** | Epha4 | 2929 | 15 | 40 |
| **Dkk1** | Lrp5 | 1 | 2948 | 41 |
| **Dlk1** | Notch3 | 149 | 2806 | 42 |
| **Lama2** | Itga6 | 2775 | 188 | 43 |
| **Efna2** | Epha1 | 1615 | 1433 | 44 |
| **Sema6a** | Plxna4 | 1619 | 1457 | 45 |
| **Cntn4** | Ptprg | 702 | 2382 | 46 |
| **Tfpi** | Vldlr | 2179 | 941 | 47 |
| **Efna3** | Epha1 | 1710 | 1433 | 48 |
| **Gm11808** | Bmpr1b | 965 | 2234 | 49 |
| **Fgf2** | Fgfrl1 | 353 | 2857 | 50 |
| **Ntn1** | Unc5b | 211 | 3030 | 51 |
| **Gnai2** | Adora1 | 743 | 2531 | 52 |
| **Col5a2** | Ddr1 | 526 | 2830 | 53 |
| **Wnt5a** | Fzd4 | 2830 | 532 | 54 |
| **Col11a1** | Ddr1 | 557 | 2830 | 55 |
| **Efnb3** | Ephb3 | 2929 | 505 | 56 |
| **Hgf** | Met | 1795 | 1677 | 57 |
| **Reln** | Itga3 | 35 | 3453 | 58 |
| **Gnai2** | Igf1r | 743 | 2775 | 59 |
| **Wnt5a** | Fzd8 | 2830 | 778 | 60 |
| **Efna4** | Epha1 | 2184 | 1433 | 61 |
| **Gnai2** | Unc5b | 743 | 3030 | 62 |
| **Tnc** | Sdc1 | 2907 | 953 | 63 |
| **Col3a1** | Ddr1 | 1055 | 2830 | 64 |
| **Col5a2** | Itga1 | 526 | 3428 | 65 |
| **Col1a1** | Ddr1 | 1140 | 2830 | 66 |
| **a** | F11r | 2047 | 1954 | 67 |
| **Col8a1** | Itga1 | 608 | 3428 | 68 |
| **Tfpi** | Sdc4 | 2179 | 1887 | 69 |
| **Col4a4** | Itga1 | 747 | 3428 | 70 |
| **Fgf11** | Fgfr4 | 3248 | 1016 | 71 |
| **Dcn** | Erbb4 | 3861 | 412 | 72 |
| **Sema3a** | Nrp1 | 217 | 4131 | 73 |
| **Fgf11** | Fgfr1 | 3248 | 1211 | 74 |
| **Fgf2** | Nrp1 | 353 | 4131 | 75 |
| **Col1a1** | Itga1 | 1140 | 3428 | 76 |
| **Col1a2** | Itga1 | 1202 | 3428 | 77 |
| **Nid1** | Itga3 | 1216 | 3453 | 78 |
| **Thbs2** | Itga6 | 4504 | 188 | 79 |
| **Wnt5a** | Ryk | 2830 | 1931 | 80 |
| **Tnc** | Sdc4 | 2907 | 1887 | 81 |
| **Ngf** | Sort1 | 1125 | 3680 | 82 |
| **Tnfsf12** | Tnfrsf12a | 2450 | 2403 | 83 |
| **Efna5** | Epha4 | 4997 | 15 | 84 |
| **Lama4** | Itga3 | 1599 | 3453 | 85 |
| **Efna3** | Ephb1 | 1710 | 3386 | 86 |
| **Fn1** | Itga3 | 1677 | 3453 | 87 |
| **Col5a1** | Itga1 | 1743 | 3428 | 88 |
| **Col6a3** | Itga1 | 1751 | 3428 | 89 |
| **Gm11808** | Erbb2 | 965 | 4262 | 90 |
| **Adm** | Ramp2 | 1475 | 3867 | 91 |
| **Mfap5** | Notch1 | 4182 | 1182 | 92 |
| **Dll3** | Notch1 | 4222 | 1182 | 93 |
| **Efna5** | Epha7 | 4997 | 409 | 94 |
| **Bmp4** | Acvr2b | 4533 | 887 | 95 |
| **Dusp18** | Itga6 | 5237 | 188 | 96 |
| **Slit2** | Gpc1 | 1772 | 3654 | 97 |
| **Dcn** | Met | 3861 | 1677 | 98 |
| **Col6a2** | Itga1 | 2153 | 3428 | 99 |
| **Col6a1** | Itga1 | 2165 | 3428 | 100 |

Supplemental Table 1 SP2-UB

| Ligand | Receptor | Ligand rank | Receptor rank | Interaction rank |
|---|---|---|---|---|
| **Gdnf** | Gfra1 | 11 | 2 | 1 |
| **Gdnf** | Ret | 11 | 3 | 2 |
| **Reln** | Vldlr | 35 | 29 | 3 |
| **Ntn1** | Neo1 | 211 | 215 | 4 |
| **Fgf2** | Fgfr2 | 353 | 137 | 5 |
| **Rspo3** | Lgr5 | 493 | 298 | 6 |
| **Gnai2** | Adora1 | 743 | 214 | 7 |
| **Cxcl12** | Ackr3 | 90 | 920 | 8 |
| **Gm11808** | Bmpr1b | 965 | 130 | 9 |
| **Ntn1** | Unc5b | 211 | 1104 | 10 |
| **Cxcl12** | Sdc4 | 90 | 1432 | 11 |
| **Nid1** | Ptprf | 1216 | 352 | 12 |
| **Vegfb** | Ret | 1671 | 3 | 13 |
| **Fgf2** | Sdc4 | 353 | 1432 | 14 |
| **Gnai2** | Unc5b | 743 | 1104 | 15 |
| **Rspo3** | Sdc4 | 493 | 1432 | 16 |
| **Gm11808** | Erbb2 | 965 | 1009 | 17 |
| **Gnai2** | Igf1r | 743 | 1258 | 18 |
| **Hgf** | St14 | 1795 | 255 | 19 |
| **Tfpi** | Vldlr | 2179 | 29 | 20 |
| **Cxcl12** | Cxcr4 | 90 | 2294 | 21 |
| **Ngf** | Sort1 | 1125 | 1356 | 22 |
| **Tbc1d24** | Neo1 | 2496 | 215 | 23 |
| **Hgf** | Met | 1795 | 1018 | 24 |
| **Fgf2** | Fgfrl1 | 353 | 2614 | 25 |
| **Ngf** | Maged1 | 1125 | 1863 | 26 |
| **Efna2** | Epha4 | 1615 | 1380 | 27 |
| **Dlk1** | Notch1 | 149 | 2893 | 28 |
| **Cntn4** | Ptprg | 702 | 2348 | 29 |
| **Efna3** | Epha4 | 1710 | 1380 | 30 |
| **Mfap2** | Notch1 | 411 | 2893 | 31 |
| **Dkk1** | Lrp6 | 1 | 3312 | 32 |
| **a** | F11r | 2047 | 1329 | 33 |
| **Fgf11** | Fgfr2 | 3248 | 137 | 34 |
| **Dkk2** | Lrp6 | 143 | 3312 | 35 |
| **Lama4** | Itga6 | 1599 | 1961 | 36 |
| **Efna4** | Epha4 | 2184 | 1380 | 37 |
| **Tfpi** | Sdc4 | 2179 | 1432 | 38 |
| **Fn1** | Itga6 | 1677 | 1961 | 39 |
| **Rspo3** | Lrp6 | 493 | 3312 | 40 |
| **C1qtnf1** | Avpr2 | 2152 | 1671 | 41 |
| **Gm11808** | Notch1 | 965 | 2893 | 42 |
| **Rgma** | Bmpr1b | 3740 | 130 | 43 |
| **Edil3** | Itgb5 | 2081 | 1810 | 44 |
| **Dkk1** | Lrp5 | 1 | 3923 | 45 |
| **Rgma** | Neo1 | 3740 | 215 | 46 |
| **Reln** | Itga3 | 35 | 3958 | 47 |
| **Efna2** | Epha1 | 1615 | 2389 | 48 |
| **Wnt5a** | Fzd4 | 2830 | 1222 | 49 |
| **Efna3** | Epha1 | 1710 | 2389 | 50 |
| **Col6a1** | Itga6 | 2165 | 1961 | 51 |
| **Efnb3** | Ephb2 | 2929 | 1248 | 52 |
| **Gdf11** | Acvr2a | 752 | 3452 | 53 |
| **Efnb3** | Epha4 | 2929 | 1380 | 54 |
| **Tnc** | Sdc4 | 2907 | 1432 | 55 |
| **Sema3a** | Plxna3 | 217 | 4314 | 56 |
| **Efna4** | Epha1 | 2184 | 2389 | 57 |
| **Tfpi** | F3 | 2179 | 2452 | 58 |
| **Igf2** | Igf1r | 3381 | 1258 | 59 |
| **Bmp4** | Bmpr1b | 4533 | 130 | 60 |
| **Lama2** | Itga6 | 2775 | 1961 | 61 |
| **Fgf2** | Sdc1 | 353 | 4403 | 62 |
| **Ntn1** | Unc5c | 211 | 4592 | 63 |
| **Sema3a** | Plxna1 | 217 | 4622 | 64 |
| **Dcn** | Met | 3861 | 1018 | 65 |
| **Igfbp4** | Lrp6 | 1633 | 3312 | 66 |
| **Fn1** | Cd79a | 1677 | 3334 | 67 |
| **Tnc** | Ptprz1 | 2907 | 2170 | 68 |
| **Nid1** | Itga3 | 1216 | 3958 | 69 |
| **Tgfb3** | Tgfbr1 | 238 | 4970 | 70 |
| **Ltbp3** | Itgb5 | 3458 | 1810 | 71 |
| **Fgf13** | Scn8a | 5451 | 63 | 72 |
| **Igf1** | Igf1r | 4288 | 1258 | 73 |
| **Lama4** | Itga3 | 1599 | 3958 | 74 |
| **Fgf13** | Fgfr2 | 5451 | 137 | 75 |
| **Fn1** | Itga3 | 1677 | 3958 | 76 |
| **Nts** | Sort1 | 4296 | 1356 | 77 |
| **Fbn1** | Itgb6 | 1363 | 4308 | 78 |
| **Tgfb2** | Tgfbr1 | 746 | 4970 | 79 |
| **Fgf2** | Fgfr4 | 353 | 5365 | 80 |
| **Slit2** | Gpc1 | 1772 | 3959 | 81 |
| **Sfrp1** | Fzd6 | 1036 | 4716 | 82 |
| **Vcam1** | Itga4 | 682 | 5129 | 83 |
| **Gm11808** | Tgfbr1 | 965 | 4970 | 84 |
| **Wnt5a** | Ror1 | 2830 | 3115 | 85 |
| **Sema3a** | Nrp1 | 217 | 5741 | 86 |
| **Itgb3bp** | Itgb5 | 4172 | 1810 | 87 |
| **Fn1** | Itgb6 | 1677 | 4308 | 88 |
| **Efnb3** | Rhbdl2 | 2929 | 3092 | 89 |
| **App** | Slc45a3 | 4360 | 1721 | 90 |
| **Fgf2** | Nrp1 | 353 | 5741 | 91 |
| **Slit2** | Sdc1 | 1772 | 4403 | 92 |
| **Hgf** | Sdc1 | 1795 | 4403 | 93 |
| **Efna5** | Ephb2 | 4997 | 1248 | 94 |
| **Col5a2** | Ddr1 | 526 | 5760 | 95 |
| **Col11a1** | Ddr1 | 557 | 5760 | 96 |
| **Efna3** | Ephb6 | 1710 | 4629 | 97 |
| **Efna5** | Epha4 | 4997 | 1380 | 98 |
| **Thbs2** | Itga6 | 4504 | 1961 | 99 |
| **Tgfb3** | Eng | 238 | 6245 | 100 |

Supplemental Table 1 SP3-RV

| Ligand | Receptor | Ligand rank | Receptor rank | Interaction rank |
|---|---|---|---|---|
| **Gdnf** | Gfra1 | 1 | 42 | 1 |
| **Rspo3** | Lgr5 | 21 | 87 | 2 |
| **Lama4** | Itga6 | 200 | 188 | 3 |
| **Efna5** | Epha4 | 389 | 15 | 4 |
| **Efna3** | Epha4 | 480 | 15 | 5 |
| **Efna4** | Epha4 | 610 | 15 | 6 |
| **Rspo3** | Lrp6 | 21 | 670 | 7 |
| **Efna5** | Epha7 | 389 | 409 | 8 |
| **Rspo3** | Fzd8 | 21 | 778 | 9 |
| **Dkk1** | Lrp6 | 182 | 670 | 10 |
| **Efna3** | Epha7 | 480 | 409 | 11 |
| **Reln** | Vldlr | 15 | 941 | 12 |
| **Hgf** | St14 | 122 | 883 | 13 |
| **Fgf2** | Sdc1 | 62 | 953 | 14 |
| **Efna4** | Epha7 | 610 | 409 | 15 |
| **Hgf** | Sdc1 | 122 | 953 | 16 |
| **Fgf2** | Fgfr4 | 62 | 1016 | 17 |
| **Fn1** | Itga6 | 965 | 188 | 18 |
| **Fgf2** | Fgfr1 | 62 | 1211 | 19 |
| **Mfap2** | Notch1 | 144 | 1182 | 20 |
| **Rspo3** | Lgr4 | 21 | 1322 | 21 |
| **Fgf2** | Gpc4 | 62 | 1359 | 22 |
| **Dlk1** | Notch1 | 322 | 1182 | 23 |
| **Gm11808** | Notch1 | 394 | 1182 | 24 |
| **Tfpi** | Vldlr | 746 | 941 | 25 |
| **Hgf** | Met | 122 | 1677 | 26 |
| **Efna5** | Epha1 | 389 | 1433 | 27 |
| **Rspo3** | Sdc4 | 21 | 1887 | 28 |
| **Efna3** | Epha1 | 480 | 1433 | 29 |
| **Fgf2** | Sdc4 | 62 | 1887 | 30 |
| **Efna4** | Epha1 | 610 | 1433 | 31 |
| **Cxcl12** | Sdc4 | 191 | 1887 | 32 |
| **Sema6a** | Plxna4 | 623 | 1457 | 33 |
| **Sema3a** | Plxna4 | 678 | 1457 | 34 |
| **Dkk2** | Lrp6 | 1487 | 670 | 35 |
| **Fgf2** | Fgfr2 | 62 | 2386 | 36 |
| **Cxcl12** | Cxcr4 | 191 | 2292 | 37 |
| **Gm11808** | Bmpr1b | 394 | 2234 | 38 |
| **Tfpi** | Sdc4 | 746 | 1887 | 39 |
| **Rgma** | Bmpr1b | 472 | 2234 | 40 |
| **a** | F11r | 847 | 1954 | 41 |
| **Fgf2** | Fgfrl1 | 62 | 2857 | 42 |
| **Fgf13** | Scn8a | 2407 | 565 | 43 |
| **Ntn1** | Unc5b | 20 | 3030 | 44 |
| **Dlk1** | Notch3 | 322 | 2806 | 45 |
| **Dkk1** | Lrp5 | 182 | 2948 | 46 |
| **Bmp5** | Acvr2b | 2263 | 887 | 47 |
| **Col11a1** | Ddr1 | 339 | 2830 | 48 |
| **Fgf13** | Fgfr4 | 2407 | 1016 | 49 |
| **Igf2** | Igf1r | 655 | 2775 | 50 |
| **Wnt5a** | Fzd4 | 2916 | 532 | 51 |
| **Reln** | Itga3 | 15 | 3453 | 52 |
| **Col4a4** | Itga1 | 73 | 3428 | 53 |
| **Col1a1** | Ddr1 | 679 | 2830 | 54 |
| **Nrtn** | Gfra1 | 3492 | 42 | 55 |
| **Col8a1** | Itga1 | 123 | 3428 | 56 |
| **Efnb1** | Epha4 | 3587 | 15 | 57 |
| **Fgf13** | Fgfr1 | 2407 | 1211 | 58 |
| **Lama4** | Itga3 | 200 | 3453 | 59 |
| **Wnt5a** | Fzd8 | 2916 | 778 | 60 |
| **Cntn4** | Ptprg | 1330 | 2382 | 61 |
| **Efna5** | Ephb1 | 389 | 3386 | 62 |
| **Efna3** | Ephb1 | 480 | 3386 | 63 |
| **Col5a2** | Ddr1 | 1184 | 2830 | 64 |
| **Tnfsf12** | Tnfrsf12a | 1624 | 2403 | 65 |
| **Efnb1** | Ephb3 | 3587 | 505 | 66 |
| **Col1a1** | Itga1 | 679 | 3428 | 67 |
| **Col4a3** | Itga1 | 721 | 3428 | 68 |
| **Efna5** | Ephb2 | 389 | 3777 | 69 |
| **Fgf2** | Nrp1 | 62 | 4131 | 70 |
| **Col3a1** | Ddr1 | 1386 | 2830 | 71 |
| **Efnb3** | Epha4 | 4330 | 15 | 72 |
| **Efna2** | Epha4 | 4350 | 15 | 73 |
| **Nid1** | Ptprf | 3848 | 568 | 74 |
| **Tfpi** | F3 | 746 | 3671 | 75 |
| **Fn1** | Itga3 | 965 | 3453 | 76 |
| **Bmp5** | Bmpr1b | 2263 | 2234 | 77 |
| **Col5a2** | Itga1 | 1184 | 3428 | 78 |
| **Gm11808** | Erbb2 | 394 | 4262 | 79 |
| **Gnai2** | Adra2a | 2594 | 2153 | 80 |
| **Efna2** | Epha7 | 4350 | 409 | 81 |
| **a** | Mgrn1 | 847 | 3929 | 82 |
| **Fgf13** | Fgfr2 | 2407 | 2386 | 83 |
| **Sema3a** | Nrp1 | 678 | 4131 | 84 |
| **Efnb3** | Ephb3 | 4330 | 505 | 85 |
| **Wnt5a** | Ryk | 2916 | 1931 | 86 |
| **Igfbp4** | Lrp6 | 4241 | 670 | 87 |
| **Col1a2** | Itga1 | 1518 | 3428 | 88 |
| **Sema3d** | Nrp1 | 823 | 4131 | 89 |
| **Col6a1** | Itga6 | 4829 | 188 | 90 |
| **Igfbp4** | Fzd8 | 4241 | 778 | 91 |
| **Dusp18** | Itga6 | 4891 | 188 | 92 |
| **Gnai2** | Adora1 | 2594 | 2531 | 93 |
| **Col6a3** | Itga1 | 1818 | 3428 | 94 |
| **Fgf11** | Fgfr4 | 4322 | 1016 | 95 |
| **Gnai2** | Igf1r | 2594 | 2775 | 96 |
| **Lamc1** | Itga6 | 5223 | 188 | 97 |
| **Penk** | Oprk1 | 406 | 5117 | 98 |
| **Fgf11** | Fgfr1 | 4322 | 1211 | 99 |
| **Gnai2** | Unc5b | 2594 | 3030 | 100 |

Supplemental Table 1 SP3-UB

| Ligand | Receptor | Ligand rank | Receptor rank | Interaction rank |
|---|---|---|---|---|
| **Gdnf** | Gfra1 | 1 | 2 | 1 |
| **Gdnf** | Ret | 1 | 3 | 2 |
| **Reln** | Vldlr | 15 | 29 | 3 |
| **Fgf2** | Fgfr2 | 62 | 137 | 4 |
| **Ntn1** | Neo1 | 20 | 215 | 5 |
| **Rspo3** | Lgr5 | 21 | 298 | 6 |
| **Hgf** | St14 | 122 | 255 | 7 |
| **Gm11808** | Bmpr1b | 394 | 130 | 8 |
| **Rgma** | Bmpr1b | 472 | 130 | 9 |
| **Rgma** | Neo1 | 472 | 215 | 10 |
| **Tfpi** | Vldlr | 746 | 29 | 11 |
| **Cxcl12** | Ackr3 | 191 | 920 | 12 |
| **Ntn1** | Unc5b | 20 | 1104 | 13 |
| **Hgf** | Met | 122 | 1018 | 14 |
| **Gm11808** | Erbb2 | 394 | 1009 | 15 |
| **Rspo3** | Sdc4 | 21 | 1432 | 16 |
| **Fgf2** | Sdc4 | 62 | 1432 | 17 |
| **Cxcl12** | Sdc4 | 191 | 1432 | 18 |
| **Efna5** | Ephb2 | 389 | 1248 | 19 |
| **Efna5** | Epha4 | 389 | 1380 | 20 |
| **Efna3** | Epha4 | 480 | 1380 | 21 |
| **Igf2** | Igf1r | 655 | 1258 | 22 |
| **Efna4** | Epha4 | 610 | 1380 | 23 |
| **Lama4** | Itga6 | 200 | 1961 | 24 |
| **a** | F11r | 847 | 1329 | 25 |
| **Tfpi** | Sdc4 | 746 | 1432 | 26 |
| **Bmp5** | Bmpr1b | 2263 | 130 | 27 |
| **Fgf13** | Scn8a | 2407 | 63 | 28 |
| **Cxcl12** | Cxcr4 | 191 | 2294 | 29 |
| **Fgf13** | Fgfr2 | 2407 | 137 | 30 |
| **Fgf2** | Fgfrl1 | 62 | 2614 | 31 |
| **Efna5** | Epha1 | 389 | 2389 | 32 |
| **Gnai2** | Adora1 | 2594 | 214 | 33 |
| **Vegfb** | Ret | 2842 | 3 | 34 |
| **Efna3** | Epha1 | 480 | 2389 | 35 |
| **Tbc1d24** | Neo1 | 2676 | 215 | 36 |
| **Fn1** | Itga6 | 965 | 1961 | 37 |
| **Efna4** | Epha1 | 610 | 2389 | 38 |
| **Ltbp3** | Itgb5 | 1203 | 1810 | 39 |
| **Mfap2** | Notch1 | 144 | 2893 | 40 |
| **Tfpi** | F3 | 746 | 2452 | 41 |
| **Dlk1** | Notch1 | 322 | 2893 | 42 |
| **Gm11808** | Notch1 | 394 | 2893 | 43 |
| **Rspo3** | Lrp6 | 21 | 3312 | 44 |
| **Nrtn** | Gfra1 | 3492 | 2 | 45 |
| **Dkk1** | Lrp6 | 182 | 3312 | 46 |
| **Nrtn** | Ret | 3492 | 3 | 47 |
| **Cntn4** | Ptprg | 1330 | 2348 | 48 |
| **Gnai2** | Unc5b | 2594 | 1104 | 49 |
| **Gnai2** | Igf1r | 2594 | 1258 | 50 |
| **Reln** | Itga3 | 15 | 3958 | 51 |
| **Ngf** | Sort1 | 2635 | 1356 | 52 |
| **Dkk1** | Lrp5 | 182 | 3923 | 53 |
| **Wnt5a** | Fzd4 | 2916 | 1222 | 54 |
| **Lama4** | Itga3 | 200 | 3958 | 55 |
| **Nid1** | Ptprf | 3848 | 352 | 56 |
| **Fn1** | Cd79a | 965 | 3334 | 57 |
| **C1qtnf1** | Avpr2 | 2636 | 1671 | 58 |
| **Fgf11** | Fgfr2 | 4322 | 137 | 59 |
| **Fgf2** | Sdc1 | 62 | 4403 | 60 |
| **Ngf** | Maged1 | 2635 | 1863 | 61 |
| **Hgf** | Sdc1 | 122 | 4403 | 62 |
| **Efnb1** | Erbb2 | 3587 | 1009 | 63 |
| **Ntn1** | Unc5c | 20 | 4592 | 64 |
| **Dkk2** | Lrp6 | 1487 | 3312 | 65 |
| **Edil3** | Itgb5 | 2995 | 1810 | 66 |
| **Efnb1** | Ephb2 | 3587 | 1248 | 67 |
| **Bmp5** | Bmpr1a | 2263 | 2644 | 68 |
| **Fn1** | Itga3 | 965 | 3958 | 69 |
| **Efnb1** | Epha4 | 3587 | 1380 | 70 |
| **Sema3a** | Plxna3 | 678 | 4314 | 71 |
| **Efna5** | Ephb6 | 389 | 4629 | 72 |
| **Sfrp1** | Fzd6 | 341 | 4716 | 73 |
| **Tgfb3** | Tgfbr1 | 87 | 4970 | 74 |
| **Efna3** | Ephb6 | 480 | 4629 | 75 |
| **Fn1** | Itgb6 | 965 | 4308 | 76 |
| **Sema3a** | Plxna1 | 678 | 4622 | 77 |
| **Tgfb2** | Tgfbr1 | 386 | 4970 | 78 |
| **Gm11808** | Tgfbr1 | 394 | 4970 | 79 |
| **Fgf2** | Fgfr4 | 62 | 5365 | 80 |
| **Efnb3** | Ephb2 | 4330 | 1248 | 81 |
| **Tgm2** | Sdc4 | 4146 | 1432 | 82 |
| **Efnb3** | Epha4 | 4330 | 1380 | 83 |
| **Bmp5** | Acvr2a | 2263 | 3452 | 84 |
| **Efna2** | Epha4 | 4350 | 1380 | 85 |
| **Fgf2** | Nrp1 | 62 | 5741 | 86 |
| **App** | Slc45a3 | 4276 | 1721 | 87 |
| **Wnt5a** | Ror1 | 2916 | 3115 | 88 |
| **Fn1** | Itga4 | 965 | 5129 | 89 |
| **Col11a1** | Ddr1 | 339 | 5760 | 90 |
| **Lpl** | Vldlr | 6115 | 29 | 91 |
| **Tgfb3** | Eng | 87 | 6245 | 92 |
| **Sema3a** | Nrp1 | 678 | 5741 | 93 |
| **Col1a1** | Ddr1 | 679 | 5760 | 94 |
| **Fbn1** | Itgb6 | 2154 | 4308 | 95 |
| **Sema3d** | Nrp1 | 823 | 5741 | 96 |
| **Tgfb2** | Eng | 386 | 6245 | 97 |
| **Efna2** | Epha1 | 4350 | 2389 | 98 |
| **Col6a1** | Itga6 | 4829 | 1961 | 99 |
| **Wnt5a** | Ryk | 2916 | 3918 | 100 |

Supplemental Table 1 RV-SP1

| Ligand | Receptor | Ligand rank | Receptor rank | Interaction rank |
|---|---|---|---|---|
| **Npy** | Fap | 8 | 16 | 1 |
| **Wnt4** | Fzd2 | 9 | 70 | 2 |
| **Pdgfa** | Pdgfra | 29 | 89 | 3 |
| **Bmp4** | Bmpr1a | 345 | 203 | 4 |
| **Efnb1** | Ephb2 | 446 | 168 | 5 |
| **Vcan** | Egfr | 280 | 379 | 6 |
| **Ncam1** | Ptpra | 215 | 605 | 7 |
| **Bmp2** | Bmpr1a | 802 | 203 | 8 |
| **Plat** | Lrp1 | 593 | 545 | 9 |
| **Rspo1** | Fzd8 | 151 | 1107 | 10 |
| **Rspo1** | Lrp6 | 151 | 1188 | 11 |
| **Tgfa** | Egfr | 1041 | 379 | 12 |
| **Lama1** | Itgb8 | 1135 | 372 | 13 |
| **Vegfa** | Ephb2 | 1362 | 168 | 14 |
| **Ntf3** | Ntrk2 | 1535 | 122 | 15 |
| **Bmp7** | Bmpr1a | 1472 | 203 | 16 |
| **Efnb2** | Ephb2 | 1520 | 168 | 17 |
| **Bmp4** | Acvr1 | 345 | 1347 | 18 |
| **Vegfa** | Egfr | 1362 | 379 | 19 |
| **Lpl** | Lrp1 | 1525 | 545 | 20 |
| **Lama1** | Sdc2 | 1135 | 986 | 21 |
| **Bmp2** | Acvr1 | 802 | 1347 | 22 |
| **Vcan** | Itgb1 | 280 | 2281 | 23 |
| **Lama4** | Itgb1 | 312 | 2281 | 24 |
| **Lamc1** | Itgb1 | 327 | 2281 | 25 |
| **Bmp7** | Acvr1 | 1472 | 1347 | 26 |
| **Lpl** | Sdc1 | 1525 | 1316 | 27 |
| **Col4a1** | Itgb8 | 2644 | 372 | 28 |
| **Gnai2** | Egfr | 2696 | 379 | 29 |
| **BC055324** | Plscr4 | 2071 | 1070 | 30 |
| **Col2a1** | Itgb1 | 891 | 2281 | 31 |
| **Pdgfa** | Pdgfrb | 29 | 3165 | 32 |
| **Vtn** | Tnfrsf11b | 3061 | 251 | 33 |
| **Lamb1** | Itgb1 | 1034 | 2281 | 34 |
| **Nppc** | Npr3 | 1719 | 1603 | 35 |
| **Hras** | Sdc2 | 2346 | 986 | 36 |
| **Vegfa** | Itga9 | 1362 | 2006 | 37 |
| **Slit3** | Robo2 | 1956 | 1427 | 38 |
| **Lamc1** | Itga1 | 327 | 3084 | 39 |
| **Lama1** | Itgb1 | 1135 | 2281 | 40 |
| **Col2a1** | Itga2b | 891 | 2530 | 41 |
| **Gm11808** | Egfr | 3043 | 379 | 42 |
| **Lama4** | Itgav | 312 | 3116 | 43 |
| **Vtn** | Itgb8 | 3061 | 372 | 44 |
| **Lamc1** | Itgav | 327 | 3116 | 45 |
| **Gdf5** | Bmpr1a | 3296 | 203 | 46 |
| **Gnai2** | Tbxa2r | 2696 | 841 | 47 |
| **Tac2** | Tacr3 | 1499 | 2122 | 48 |
| **Gnai2** | Agtr2 | 2696 | 928 | 49 |
| **Hras** | Cav1 | 2346 | 1294 | 50 |
| **Vegfa** | Itgb1 | 1362 | 2281 | 51 |
| **Lamc2** | Cd151 | 2453 | 1269 | 52 |
| **Prss23** | Tmem222 | 1048 | 2805 | 53 |
| **Col4a1** | Cd47 | 2644 | 1260 | 54 |
| **Col2a1** | Itga1 | 891 | 3084 | 55 |
| **Gnai2** | Cav1 | 2696 | 1294 | 56 |
| **Sema3f** | Nrp1 | 748 | 3252 | 57 |
| **L1cam** | Ephb2 | 3935 | 168 | 58 |
| **Lamb1** | Itga1 | 1034 | 3084 | 59 |
| **Gdf5** | Ror2 | 3296 | 825 | 60 |
| **Timp2** | Itgb1 | 1841 | 2281 | 61 |
| **Cdh1** | Egfr | 3752 | 379 | 62 |
| **Lamb1** | Itgav | 1034 | 3116 | 63 |
| **Lama1** | Itga1 | 1135 | 3084 | 64 |
| **Sost** | Lrp6 | 3095 | 1188 | 65 |
| **L1cam** | Egfr | 3935 | 379 | 66 |
| **Rgmb** | Neo1 | 3588 | 726 | 67 |
| **Vtn** | Cd47 | 3061 | 1260 | 68 |
| **Cdh1** | Cdh2 | 3752 | 585 | 69 |
| **Adam10** | Axl | 3668 | 707 | 70 |
| **Gm11808** | Acvr1 | 3043 | 1347 | 71 |
| **Slit2** | Sdc1 | 3090 | 1316 | 72 |
| **Vegfa** | Itgav | 1362 | 3116 | 73 |
| **Npy** | Npy2r | 8 | 4478 | 74 |
| **Slit2** | Robo2 | 3090 | 1427 | 75 |
| **Efnb1** | Ephb3 | 446 | 4130 | 76 |
| **Col18a1** | Gpc4 | 3600 | 992 | 77 |
| **Vegfa** | Nrp1 | 1362 | 3252 | 78 |
| **Gdf5** | Acvr1 | 3296 | 1347 | 79 |
| **Sema4c** | Plxnb2 | 3580 | 1101 | 80 |
| **Lamc2** | Itgb1 | 2453 | 2281 | 81 |
| **Vegfc** | Itga9 | 2908 | 2006 | 82 |
| **Col4a1** | Itgb1 | 2644 | 2281 | 83 |
| **Efnb1** | Ephb4 | 446 | 4581 | 84 |
| **Mfge8** | Itgav | 2059 | 3116 | 85 |
| **Vegfc** | Itgb1 | 2908 | 2281 | 86 |
| **Efna5** | Ephb2 | 5025 | 168 | 87 |
| **Gnai2** | S1pr3 | 2696 | 2500 | 88 |
| **Mfge8** | Pdgfrb | 2059 | 3165 | 89 |
| **Pdgfc** | Pdgfra | 5137 | 89 | 90 |
| **Lama1** | Gpc1 | 1135 | 4138 | 91 |
| **Serpina1a** | Lrp1 | 4787 | 545 | 92 |
| **Vtn** | Itgb1 | 3061 | 2281 | 93 |
| **Vegfa** | Kdr | 1362 | 3991 | 94 |
| **Lama5** | Sdc1 | 4093 | 1316 | 95 |
| **Npnt** | Itgb1 | 3150 | 2281 | 96 |
| **Fbln1** | Itgb1 | 3199 | 2281 | 97 |
| **Vegfa** | Gpc1 | 1362 | 4138 | 98 |
| **Lama1** | Nt5e | 1135 | 4371 | 99 |
| **Clcf1** | Il6st | 1341 | 4172 | 100 |

Supplemental Table 1 RV-SP2

| Ligand | Receptor | Ligand rank | Receptor rank | Interaction rank |
|---|---|---|---|---|
| **Pdgfa** | Pdgfrb | 29 | 23 | 1 |
| **Pdgfa** | Pdgfra | 29 | 756 | 2 |
| **Sema3f** | Nrp1 | 748 | 215 | 3 |
| **Ncam1** | Fgfr1 | 215 | 888 | 4 |
| **Npy** | Fap | 8 | 1236 | 5 |
| **Vegfa** | Itga9 | 1362 | 130 | 6 |
| **Lamc1** | Itga1 | 327 | 1225 | 7 |
| **Vegfa** | Nrp1 | 1362 | 215 | 8 |
| **Vcan** | Egfr | 280 | 1299 | 9 |
| **Jag1** | Notch3 | 94 | 1521 | 10 |
| **Fgf8** | Fgfr1 | 977 | 888 | 11 |
| **Fgf15** | Fgfr1 | 1023 | 888 | 12 |
| **Mfge8** | Pdgfrb | 2059 | 23 | 13 |
| **Dll1** | Notch3 | 583 | 1521 | 14 |
| **Col2a1** | Itga1 | 891 | 1225 | 15 |
| **Ncam1** | Robo1 | 215 | 1934 | 16 |
| **Lama1** | Itgb8 | 1135 | 1100 | 17 |
| **Lamb1** | Itga1 | 1034 | 1225 | 18 |
| **Bmp2** | Eng | 802 | 1503 | 19 |
| **Tgfa** | Egfr | 1041 | 1299 | 20 |
| **Lama1** | Itga1 | 1135 | 1225 | 21 |
| **Ncam1** | Cacna1c | 215 | 2215 | 22 |
| **Vcan** | Itgb1 | 280 | 2286 | 23 |
| **Lama4** | Itgb1 | 312 | 2286 | 24 |
| **Lamc1** | Itgb1 | 327 | 2286 | 25 |
| **Jag1** | Notch1 | 94 | 2535 | 26 |
| **Wnt4** | Fzd2 | 9 | 2644 | 27 |
| **Vegfa** | Egfr | 1362 | 1299 | 28 |
| **Bmp4** | Bmpr1a | 345 | 2345 | 29 |
| **Bmp4** | Acvr1 | 345 | 2361 | 30 |
| **Gnai2** | F2r | 2696 | 59 | 31 |
| **Gnai2** | S1pr3 | 2696 | 106 | 32 |
| **Plat** | Lrp1 | 593 | 2219 | 33 |
| **Clcf1** | Cntfr | 1341 | 1550 | 34 |
| **Gnai2** | Ednra | 2696 | 208 | 35 |
| **Bmp7** | Eng | 1472 | 1503 | 36 |
| **Vegfc** | Itga9 | 2908 | 130 | 37 |
| **Gnai2** | Agtr2 | 2696 | 385 | 38 |
| **Prss23** | Tmem222 | 1048 | 2058 | 39 |
| **Dll1** | Notch1 | 583 | 2535 | 40 |
| **Bmp2** | Bmpr1a | 802 | 2345 | 41 |
| **Bmp2** | Acvr1 | 802 | 2361 | 42 |
| **Col2a1** | Itgb1 | 891 | 2286 | 43 |
| **Ltbp1** | Itgb5 | 462 | 2719 | 44 |
| **Lamb1** | Itgb1 | 1034 | 2286 | 45 |
| **Lama1** | Itgb1 | 1135 | 2286 | 46 |
| **Vtn** | Itga8 | 3061 | 521 | 47 |
| **Vegfa** | Itgb1 | 1362 | 2286 | 48 |
| **Npnt** | Itga8 | 3150 | 521 | 49 |
| **Col4a1** | Itgb8 | 2644 | 1100 | 50 |
| **Lpl** | Lrp1 | 1525 | 2219 | 51 |
| **Bmp7** | Bmpr1a | 1472 | 2345 | 52 |
| **Bmp4** | Acvr2a | 345 | 3481 | 53 |
| **Bmp7** | Acvr1 | 1472 | 2361 | 54 |
| **Col4a1** | Itga1 | 2644 | 1225 | 55 |
| **Slit3** | Robo2 | 1956 | 1917 | 56 |
| **Itgb3bp** | Itgb5 | 1180 | 2719 | 57 |
| **Gnai2** | Egfr | 2696 | 1299 | 58 |
| **Lama4** | Itgav | 312 | 3688 | 59 |
| **Lamc1** | Itgav | 327 | 3688 | 60 |
| **Fgf9** | Fgfr1 | 3166 | 888 | 61 |
| **Bmp4** | Bmpr2 | 345 | 3711 | 62 |
| **Edn1** | Ednra | 3859 | 208 | 63 |
| **Gdf5** | Ror2 | 3296 | 786 | 64 |
| **Timp2** | Itgb1 | 1841 | 2286 | 65 |
| **Vtn** | Itgb8 | 3061 | 1100 | 66 |
| **Clcf1** | Il6st | 1341 | 2891 | 67 |
| **Gnai2** | Ptpru | 2696 | 1583 | 68 |
| **Bmp2** | Acvr2a | 802 | 3481 | 69 |
| **Gm11808** | Egfr | 3043 | 1299 | 70 |
| **Nppc** | Npr3 | 1719 | 2743 | 71 |
| **Adam10** | Axl | 3668 | 824 | 72 |
| **Bmp2** | Bmpr2 | 802 | 3711 | 73 |
| **Jag1** | Notch4 | 94 | 4507 | 74 |
| **Lamb1** | Itgav | 1034 | 3688 | 75 |
| **Lamc2** | Itgb1 | 2453 | 2286 | 76 |
| **Hras** | Agtr1a | 2346 | 2400 | 77 |
| **Col4a1** | Itgb1 | 2644 | 2286 | 78 |
| **Bmp7** | Acvr2a | 1472 | 3481 | 79 |
| **Slit2** | Robo2 | 3090 | 1917 | 80 |
| **Slit2** | Robo1 | 3090 | 1934 | 81 |
| **Vegfa** | Itgav | 1362 | 3688 | 82 |
| **Cdh1** | Egfr | 3752 | 1299 | 83 |
| **Adam15** | Itga9 | 4927 | 130 | 84 |
| **Nppc** | Npr2 | 1719 | 3339 | 85 |
| **Dll1** | Notch4 | 583 | 4507 | 86 |
| **Ntf3** | Ntrk2 | 1535 | 3569 | 87 |
| **Pdgfc** | Pdgfrb | 5137 | 23 | 88 |
| **Bmp7** | Bmpr2 | 1472 | 3711 | 89 |
| **Lpl** | Sdc1 | 1525 | 3661 | 90 |
| **Sema3f** | Plxna3 | 748 | 4444 | 91 |
| **Vegfc** | Itgb1 | 2908 | 2286 | 92 |
| **L1cam** | Egfr | 3935 | 1299 | 93 |
| **Vtn** | Itgb1 | 3061 | 2286 | 94 |
| **Sema3f** | Plxna1 | 748 | 4650 | 95 |
| **Gm11808** | Acvr1 | 3043 | 2361 | 96 |
| **Npnt** | Itgb1 | 3150 | 2286 | 97 |
| **Ncam1** | Ptpra | 215 | 5221 | 98 |
| **Gm11808** | Agtr1a | 3043 | 2400 | 99 |
| **Fbln1** | Itgb1 | 3199 | 2286 | 100 |

Supplemental Table 1 RV-SP3

| Ligand | Receptor | Ligand rank | Receptor rank | Interaction rank |
|---|---|---|---|---|
| **Npy** | Fap | 8 | 363 | 1 |
| **Pdgfa** | Pdgfra | 29 | 424 | 2 |
| **Ncam1** | Robo1 | 215 | 311 | 3 |
| **Vcan** | Egfr | 280 | 439 | 4 |
| **Ncam1** | Fgfr1 | 215 | 513 | 5 |
| **Wnt4** | Fzd2 | 9 | 1160 | 6 |
| **Lama1** | Itgb8 | 1135 | 113 | 7 |
| **Pdgfa** | Pdgfrb | 29 | 1304 | 8 |
| **Efna4** | Epha7 | 784 | 597 | 9 |
| **Tgfa** | Egfr | 1041 | 439 | 10 |
| **Fgf8** | Fgfr1 | 977 | 513 | 11 |
| **Fgf15** | Fgfr1 | 1023 | 513 | 12 |
| **Bmp4** | Bmpr1a | 345 | 1247 | 13 |
| **Vegfa** | Itga9 | 1362 | 245 | 14 |
| **Vegfa** | Egfr | 1362 | 439 | 15 |
| **Bmp2** | Bmpr1a | 802 | 1247 | 16 |
| **Sema3f** | Nrp1 | 748 | 1661 | 17 |
| **Ntf3** | Ntrk2 | 1535 | 882 | 18 |
| **Slit3** | Robo2 | 1956 | 537 | 19 |
| **Ncam1** | Cacna1c | 215 | 2376 | 20 |
| **Efnb1** | Ephb2 | 446 | 2153 | 21 |
| **Bmp7** | Bmpr1a | 1472 | 1247 | 22 |
| **Col4a1** | Itgb8 | 2644 | 113 | 23 |
| **Gnai2** | F2r | 2696 | 217 | 24 |
| **Vegfa** | Nrp1 | 1362 | 1661 | 25 |
| **Efna1** | Epha7 | 2447 | 597 | 26 |
| **Gnai2** | Egfr | 2696 | 439 | 27 |
| **Vegfc** | Itga9 | 2908 | 245 | 28 |
| **Vtn** | Itgb8 | 3061 | 113 | 29 |
| **Ncam1** | Ptpra | 215 | 2986 | 30 |
| **Tac2** | Tacr3 | 1499 | 1835 | 31 |
| **Vtn** | Tnfrsf11b | 3061 | 292 | 32 |
| **Mfge8** | Pdgfrb | 2059 | 1304 | 33 |
| **Bmp2** | Eng | 802 | 2586 | 34 |
| **Slit2** | Robo1 | 3090 | 311 | 35 |
| **Gnai2** | Agtr2 | 2696 | 710 | 36 |
| **Clcf1** | Cntfr | 1341 | 2119 | 37 |
| **Gm11808** | Egfr | 3043 | 439 | 38 |
| **Vegfa** | Ephb2 | 1362 | 2153 | 39 |
| **Efna2** | Epha7 | 2978 | 597 | 40 |
| **Bmp4** | Acvr1 | 345 | 3234 | 41 |
| **Gnai2** | Ptpru | 2696 | 894 | 42 |
| **Slit2** | Robo2 | 3090 | 537 | 43 |
| **Efnb2** | Ephb2 | 1520 | 2153 | 44 |
| **Fgf9** | Fgfr1 | 3166 | 513 | 45 |
| **Sema3f** | Plxna1 | 748 | 3162 | 46 |
| **Plat** | Lrp1 | 593 | 3426 | 47 |
| **Bmp2** | Acvr1 | 802 | 3234 | 48 |
| **Bmp7** | Eng | 1472 | 2586 | 49 |
| **Efnb1** | Ephb3 | 446 | 3684 | 50 |
| **Sema3f** | Plxna3 | 748 | 3409 | 51 |
| **Ncam1** | Gfra1 | 215 | 3947 | 52 |
| **Cdh1** | Egfr | 3752 | 439 | 53 |
| **Prss23** | Tmem222 | 1048 | 3148 | 54 |
| **Gnai2** | S1pr3 | 2696 | 1647 | 55 |
| **Gdf5** | Ror2 | 3296 | 1064 | 56 |
| **L1cam** | Egfr | 3935 | 439 | 57 |
| **Efnb1** | Ephb4 | 446 | 3962 | 58 |
| **Jag1** | Notch2 | 94 | 4420 | 59 |
| **Gdf5** | Bmpr1a | 3296 | 1247 | 60 |
| **Lpl** | Sdc1 | 1525 | 3092 | 61 |
| **Bmp7** | Acvr1 | 1472 | 3234 | 62 |
| **Clec11a** | Kit | 2642 | 2151 | 63 |
| **Lpl** | Lrp1 | 1525 | 3426 | 64 |
| **Dll1** | Notch2 | 583 | 4420 | 65 |
| **Adam10** | Axl | 3668 | 1500 | 66 |
| **Adam15** | Itga9 | 4927 | 245 | 67 |
| **Jag1** | Notch1 | 94 | 5097 | 68 |
| **Efnb2** | Ephb3 | 1520 | 3684 | 69 |
| **Cdh1** | Cdh2 | 3752 | 1499 | 70 |
| **Col2a1** | Itga2b | 891 | 4457 | 71 |
| **Vcan** | Itgb1 | 280 | 5139 | 72 |
| **Rspo1** | Fzd8 | 151 | 5270 | 73 |
| **Lama4** | Itgb1 | 312 | 5139 | 74 |
| **Lamc1** | Itgb1 | 327 | 5139 | 75 |
| **Efnb2** | Ephb4 | 1520 | 3962 | 76 |
| **Bmp4** | Bmpr2 | 345 | 5145 | 77 |
| **Pdgfc** | Pdgfra | 5137 | 424 | 78 |
| **Fgf16** | Fgfr1 | 5097 | 513 | 79 |
| **Efna5** | Epha7 | 5025 | 597 | 80 |
| **Dll1** | Notch1 | 583 | 5097 | 81 |
| **Jag1** | Notch4 | 94 | 5833 | 82 |
| **Bmp2** | Bmpr2 | 802 | 5145 | 83 |
| **Fgf10** | Fgfr1 | 5465 | 513 | 84 |
| **Col2a1** | Itgb1 | 891 | 5139 | 85 |
| **Sema4a** | Plxnd1 | 4449 | 1618 | 86 |
| **L1cam** | Ephb2 | 3935 | 2153 | 87 |
| **Lamb1** | Itgb1 | 1034 | 5139 | 88 |
| **Ltbp1** | Itgb5 | 462 | 5717 | 89 |
| **Slit2** | Sdc1 | 3090 | 3092 | 90 |
| **Lama1** | Itgb1 | 1135 | 5139 | 91 |
| **Gm11808** | Acvr1 | 3043 | 3234 | 92 |
| **Nppc** | Npr2 | 1719 | 4618 | 93 |
| **Tgfa** | Erbb4 | 1041 | 5315 | 94 |
| **Rspo1** | Lrp6 | 151 | 6225 | 95 |
| **Dll1** | Notch4 | 583 | 5833 | 96 |
| **Pdgfc** | Pdgfrb | 5137 | 1304 | 97 |
| **Nppc** | Npr3 | 1719 | 4756 | 98 |
| **Vegfa** | Itgb1 | 1362 | 5139 | 99 |
| **Gdf5** | Acvr1 | 3296 | 3234 | 100 |

Supplemental Table 1 RV-UB

| Ligand | Receptor | Ligand rank | Receptor rank | Interaction rank |
|---|---|---|---|---|
| **Ncam1** | Gfra1 | 215 | 2 | 1 |
| **Ncam1** | Fgfr2 | 215 | 137 | 2 |
| **Rspo1** | Lgr5 | 151 | 298 | 3 |
| **Bmp4** | Bmpr1b | 345 | 130 | 4 |
| **Bmp2** | Bmpr1b | 802 | 130 | 5 |
| **Fgf8** | Fgfr2 | 977 | 137 | 6 |
| **Fgf15** | Fgfr2 | 1023 | 137 | 7 |
| **Clcf1** | Crlf1 | 1341 | 10 | 8 |
| **Vegfa** | Ret | 1362 | 3 | 9 |
| **Efnb1** | Erbb2 | 446 | 1009 | 10 |
| **Lpl** | Vldlr | 1525 | 29 | 11 |
| **Bmp7** | Bmpr1b | 1472 | 130 | 12 |
| **Pcsk9** | Vldlr | 1583 | 29 | 13 |
| **Efnb1** | Ephb2 | 446 | 1248 | 14 |
| **Efnb1** | Epha4 | 446 | 1380 | 15 |
| **Rspo1** | Znrf3 | 151 | 1829 | 16 |
| **Tgfa** | Erbb2 | 1041 | 1009 | 17 |
| **Efna4** | Epha4 | 784 | 1380 | 18 |
| **Ltbp1** | Itgb5 | 462 | 1810 | 19 |
| **Lama4** | Itga6 | 312 | 1961 | 20 |
| **Lamc1** | Itga6 | 327 | 1961 | 21 |
| **Lama1** | Sdc4 | 1135 | 1432 | 22 |
| **Vegfa** | Ephb2 | 1362 | 1248 | 23 |
| **Rspo4** | Lgr5 | 2366 | 298 | 24 |
| **Efnb2** | Ephb2 | 1520 | 1248 | 25 |
| **Tgfa** | Erbb3 | 1041 | 1843 | 26 |
| **Efnb2** | Epha4 | 1520 | 1380 | 27 |
| **Gnai2** | Adora1 | 2696 | 214 | 28 |
| **Jag1** | Notch1 | 94 | 2893 | 29 |
| **Bmp4** | Bmpr1a | 345 | 2644 | 30 |
| **Itgb3bp** | Itgb5 | 1180 | 1810 | 31 |
| **Lamb1** | Itga6 | 1034 | 1961 | 32 |
| **Lama1** | Itga6 | 1135 | 1961 | 33 |
| **Gm11808** | Bmpr1b | 3043 | 130 | 34 |
| **Efna4** | Epha1 | 784 | 2389 | 35 |
| **Fgf9** | Fgfr2 | 3166 | 137 | 36 |
| **Gdf5** | Bmpr1b | 3296 | 130 | 37 |
| **Bmp2** | Bmpr1a | 802 | 2644 | 38 |
| **Rspo1** | Lrp6 | 151 | 3312 | 39 |
| **Dll1** | Notch1 | 583 | 2893 | 40 |
| **Rgmb** | Bmpr1b | 3588 | 130 | 41 |
| **Bmp4** | Acvr2a | 345 | 3452 | 42 |
| **Gnai2** | Unc5b | 2696 | 1104 | 43 |
| **Rgmb** | Neo1 | 3588 | 215 | 44 |
| **Efna1** | Epha4 | 2447 | 1380 | 45 |
| **Sema5a** | Met | 2866 | 1018 | 46 |
| **Gnai2** | Igf1r | 2696 | 1258 | 47 |
| **Gm11808** | Erbb2 | 3043 | 1009 | 48 |
| **L1cam** | Fgfr2 | 3935 | 137 | 49 |
| **Spint1** | St14 | 3848 | 255 | 50 |
| **Cdh1** | Ptprf | 3752 | 352 | 51 |
| **Bmp7** | Bmpr1a | 1472 | 2644 | 52 |
| **Igf2** | Igf1r | 2986 | 1258 | 53 |
| **Bmp2** | Acvr2a | 802 | 3452 | 54 |
| **Lamc1** | Itgb4 | 327 | 3929 | 55 |
| **Lama4** | Itga3 | 312 | 3958 | 56 |
| **Lamc1** | Itga3 | 327 | 3958 | 57 |
| **Efna2** | Epha4 | 2978 | 1380 | 58 |
| **Lamc2** | Itga6 | 2453 | 1961 | 59 |
| **Efnb2** | Rhbdl2 | 1520 | 3092 | 60 |
| **Wnt4** | Fzd6 | 9 | 4716 | 61 |
| **Lamc2** | Cd151 | 2453 | 2277 | 62 |
| **Efna1** | Epha1 | 2447 | 2389 | 63 |
| **Lama5** | Bcam | 4093 | 772 | 64 |
| **Vtn** | Itgb5 | 3061 | 1810 | 65 |
| **Bmp7** | Acvr2a | 1472 | 3452 | 66 |
| **Col2a1** | Itga10 | 891 | 4048 | 67 |
| **L1cam** | Erbb2 | 3935 | 1009 | 68 |
| **Agrn** | Lrp4 | 2597 | 2356 | 69 |
| **Lamb1** | Itgb4 | 1034 | 3929 | 70 |
| **Lamb1** | Itga3 | 1034 | 3958 | 71 |
| **Cdh1** | Igf1r | 3752 | 1258 | 72 |
| **Tgfa** | Erbb4 | 1041 | 3994 | 73 |
| **Sema3f** | Plxna3 | 748 | 4314 | 74 |
| **Lama1** | Itgb4 | 1135 | 3929 | 75 |
| **Efnb1** | Ephb6 | 446 | 4629 | 76 |
| **Lama1** | Itga3 | 1135 | 3958 | 77 |
| **Lama1** | Gpc1 | 1135 | 3959 | 78 |
| **L1cam** | Ephb2 | 3935 | 1248 | 79 |
| **Fgf16** | Fgfr2 | 5097 | 137 | 80 |
| **Vegfa** | Gpc1 | 1362 | 3959 | 81 |
| **Efna2** | Epha1 | 2978 | 2389 | 82 |
| **Sema3f** | Plxna1 | 748 | 4622 | 83 |
| **Efnb1** | Ephb3 | 446 | 4939 | 84 |
| **Vcan** | Itga4 | 280 | 5129 | 85 |
| **Lamc3** | Itga6 | 3462 | 1961 | 86 |
| **Vegfa** | Sirpa | 1362 | 4078 | 87 |
| **Prss23** | Tmem222 | 1048 | 4396 | 88 |
| **Cdh1** | Erbb3 | 3752 | 1843 | 89 |
| **Fgf10** | Fgfr2 | 5465 | 137 | 90 |
| **Cthrc1** | Fzd3 | 3806 | 1947 | 91 |
| **L1cam** | Erbb3 | 3935 | 1843 | 92 |
| **Timp2** | Itga3 | 1841 | 3958 | 93 |
| **Lpl** | Sdc1 | 1525 | 4403 | 94 |
| **Gm11808** | Notch1 | 3043 | 2893 | 95 |
| **Gdf5** | Bmpr1a | 3296 | 2644 | 96 |
| **Npy** | Npy2r | 8 | 6038 | 97 |
| **Lama5** | Itga6 | 4093 | 1961 | 98 |
| **Sema4d** | Erbb2 | 5069 | 1009 | 99 |
| **Sema4d** | Met | 5069 | 1018 | 100 |

Supplemental Table 1 UB-SP1

| Ligand | Receptor | Ligand rank | Receptor rank | Interaction rank |
|---|---|---|---|---|
| **Nrtn** | Gfra2 | 17 | 124 | 1 |
| **Bmp7** | Bmpr1a | 59 | 203 | 2 |
| **Cdh1** | Egfr | 261 | 379 | 3 |
| **Btc** | Egfr | 322 | 379 | 4 |
| **Cdh1** | Cdh2 | 261 | 585 | 5 |
| **L1cam** | Ephb2 | 705 | 168 | 6 |
| **Npy** | Fap | 1017 | 16 | 7 |
| **L1cam** | Egfr | 705 | 379 | 8 |
| **Wnt11** | Fzd7 | 41 | 1080 | 9 |
| **Bmp7** | Acvr1 | 59 | 1347 | 10 |
| **Lamc2** | Cd151 | 172 | 1269 | 11 |
| **Slit2** | Sdc1 | 139 | 1316 | 12 |
| **Slit2** | Robo2 | 139 | 1427 | 13 |
| **Pdgfc** | Pdgfra | 1664 | 89 | 14 |
| **Sema6a** | Plxna2 | 92 | 1959 | 15 |
| **Efnb2** | Ephb2 | 2019 | 168 | 16 |
| **Lama5** | Sdc1 | 1057 | 1316 | 17 |
| **Lamc2** | Itgb1 | 172 | 2281 | 18 |
| **Col18a1** | Gpc4 | 1610 | 992 | 19 |
| **Npnt** | Itgb1 | 338 | 2281 | 20 |
| **Cgn** | F11r | 2929 | 295 | 21 |
| **Lama5** | Itgb1 | 1057 | 2281 | 22 |
| **Pdgfa** | Pdgfra | 3390 | 89 | 23 |
| **Hspg2** | Ptprs | 2912 | 764 | 24 |
| **Lamc1** | Itgb1 | 1441 | 2281 | 25 |
| **L1cam** | Itgav | 705 | 3116 | 26 |
| **Col18a1** | Itgb1 | 1610 | 2281 | 27 |
| **Bmp8b** | Bmpr1a | 3723 | 203 | 28 |
| **Lamb1** | Itgb1 | 1668 | 2281 | 29 |
| **Sema3d** | Nrp1 | 749 | 3252 | 30 |
| **Sema3c** | Nrp1 | 771 | 3252 | 31 |
| **BC055324** | Plscr4 | 2963 | 1070 | 32 |
| **Hspg2** | Sdc1 | 2912 | 1316 | 33 |
| **Shh** | Boc | 4144 | 114 | 34 |
| **Slit2** | Gpc1 | 139 | 4138 | 35 |
| **Col4a1** | Itgb8 | 3909 | 372 | 36 |
| **Sema3f** | Nrp1 | 1051 | 3252 | 37 |
| **Gm11808** | Egfr | 3949 | 379 | 38 |
| **Shh** | Cdon | 4144 | 217 | 39 |
| **Prss23** | Tmem222 | 1609 | 2805 | 40 |
| **Lamc1** | Itga1 | 1441 | 3084 | 41 |
| **Ctf1** | Il6st | 379 | 4172 | 42 |
| **Lamc1** | Itgav | 1441 | 3116 | 43 |
| **Lamb1** | Itga1 | 1668 | 3084 | 44 |
| **Lamb1** | Itgav | 1668 | 3116 | 45 |
| **Shh** | Smo | 4144 | 681 | 46 |
| **Pdgfc** | Pdgfrb | 1664 | 3165 | 47 |
| **Serpine2** | Lrp1 | 4321 | 545 | 48 |
| **Tac1** | Tacr3 | 2816 | 2122 | 49 |
| **App** | Lrp1 | 4431 | 545 | 50 |
| **Bmp8b** | Acvr1 | 3723 | 1347 | 51 |
| **Col4a1** | Cd47 | 3909 | 1260 | 52 |
| **Ihh** | Boc | 5074 | 114 | 53 |
| **Hspg2** | Itgb1 | 2912 | 2281 | 54 |
| **Sema4d** | Plxnb2 | 4111 | 1101 | 55 |
| **Ihh** | Cdon | 5074 | 217 | 56 |
| **Gm11808** | Acvr1 | 3949 | 1347 | 57 |
| **Shh** | Hhip | 4144 | 1195 | 58 |
| **Bcan** | Egfr | 5100 | 379 | 59 |
| **Npy** | Npy2r | 1017 | 4478 | 60 |
| **Col9a3** | Mag | 38 | 5458 | 61 |
| **Efna5** | Ephb2 | 5351 | 168 | 62 |
| **Gdf5** | Bmpr1a | 5373 | 203 | 63 |
| **Lama1** | Itgb8 | 5217 | 372 | 64 |
| **Col18a1** | Kdr | 1610 | 3991 | 65 |
| **Pdgfc** | Kdr | 1664 | 3991 | 66 |
| **App** | Cav1 | 4431 | 1294 | 67 |
| **Col18a1** | Gpc1 | 1610 | 4138 | 68 |
| **Efna4** | Epha5 | 53 | 5735 | 69 |
| **Wnt5a** | Fzd2 | 5788 | 70 | 70 |
| **Tgfa** | Egfr | 5582 | 379 | 71 |
| **Btc** | Erbb2 | 322 | 5746 | 72 |
| **Efnb2** | Ephb3 | 2019 | 4130 | 73 |
| **Col4a1** | Itgb1 | 3909 | 2281 | 74 |
| **Gdf5** | Ror2 | 5373 | 825 | 75 |
| **Lama1** | Sdc2 | 5217 | 986 | 76 |
| **Wnt4** | Fzd2 | 6163 | 70 | 77 |
| **Cgn** | Tgfbr2 | 2929 | 3323 | 78 |
| **1700013F07Rik** | Plscr4 | 5189 | 1070 | 79 |
| **Ihh** | Hhip | 5074 | 1195 | 80 |
| **Sema3c** | Nrp2 | 771 | 5501 | 81 |
| **Serpinc1** | Lrp1 | 5752 | 545 | 82 |
| **Lamc2** | Itga3 | 172 | 6141 | 83 |
| **Vegfa** | Ephb2 | 6167 | 168 | 84 |
| **L1cam** | Erbb2 | 705 | 5746 | 85 |
| **Bmp3** | Bmpr1a | 6286 | 203 | 86 |
| **Col9a1** | Mag | 1031 | 5458 | 87 |
| **Vegfa** | Egfr | 6167 | 379 | 88 |
| **Sema3f** | Nrp2 | 1051 | 5501 | 89 |
| **Pdgfa** | Pdgfrb | 3390 | 3165 | 90 |
| **Efnb2** | Ephb4 | 2019 | 4581 | 91 |
| **Shh** | Ptch2 | 4144 | 2457 | 92 |
| **Wnt5a** | Ror2 | 5788 | 825 | 93 |
| **Il11** | Gm13305 | 6455 | 162 | 94 |
| **Anxa1** | Egfr | 6255 | 379 | 95 |
| **Lamb3** | Cd151 | 5432 | 1269 | 96 |
| **Gdf5** | Acvr1 | 5373 | 1347 | 97 |
| **Serpinc1** | Sdc2 | 5752 | 986 | 98 |
| **Lrpap1** | Lrp1 | 6231 | 545 | 99 |
| **Dusp18** | Cd151 | 5562 | 1269 | 100 |

Supplemental Table 1 UB-SP2

| Ligand | Receptor | Ligand rank | Receptor rank | Interaction rank |
|---|---|---|---|---|
| **Kitl** | Epor | 114 | 378 | 1 |
| **Npnt** | Itga8 | 338 | 521 | 2 |
| **Sema3d** | Nrp1 | 749 | 215 | 3 |
| **Sema3c** | Nrp1 | 771 | 215 | 4 |
| **Sema3f** | Nrp1 | 1051 | 215 | 5 |
| **Cdh1** | Egfr | 261 | 1299 | 6 |
| **Bmp7** | Eng | 59 | 1503 | 7 |
| **Btc** | Egfr | 322 | 1299 | 8 |
| **Pdgfc** | Pdgfrb | 1664 | 23 | 9 |
| **L1cam** | Egfr | 705 | 1299 | 10 |
| **Fgf9** | Fgfr1 | 1118 | 888 | 11 |
| **Wnt11** | Fzd7 | 41 | 1999 | 12 |
| **Slit2** | Robo2 | 139 | 1917 | 13 |
| **Slit2** | Robo1 | 139 | 1934 | 14 |
| **Npy** | Fap | 1017 | 1236 | 15 |
| **Bmp7** | Bmpr1a | 59 | 2345 | 16 |
| **Bmp7** | Acvr1 | 59 | 2361 | 17 |
| **Pdgfc** | Pdgfra | 1664 | 756 | 18 |
| **Lamc2** | Itgb1 | 172 | 2286 | 19 |
| **Npnt** | Itgb1 | 338 | 2286 | 20 |
| **Lamc1** | Itga1 | 1441 | 1225 | 21 |
| **Fgf12** | Fgfr1 | 1951 | 888 | 22 |
| **Lamb1** | Itga1 | 1668 | 1225 | 23 |
| **Ctf1** | Il6st | 379 | 2891 | 24 |
| **Jag1** | Notch3 | 1811 | 1521 | 25 |
| **Lama5** | Itgb1 | 1057 | 2286 | 26 |
| **Sema3c** | Plxnd1 | 771 | 2573 | 27 |
| **Pdgfa** | Pdgfrb | 3390 | 23 | 28 |
| **Bmp7** | Acvr2a | 59 | 3481 | 29 |
| **Prss23** | Tmem222 | 1609 | 2058 | 30 |
| **Lamc1** | Itgb1 | 1441 | 2286 | 31 |
| **Bmp7** | Bmpr2 | 59 | 3711 | 32 |
| **Slit2** | Sdc1 | 139 | 3661 | 33 |
| **Cntn2** | Nrp1 | 3656 | 215 | 34 |
| **Col18a1** | Itgb1 | 1610 | 2286 | 35 |
| **Lamb1** | Itgb1 | 1668 | 2286 | 36 |
| **Pdgfa** | Pdgfra | 3390 | 756 | 37 |
| **Wnt7b** | Fzd1 | 3770 | 416 | 38 |
| **Itgb3bp** | Itgb5 | 1539 | 2719 | 39 |
| **Shh** | Boc | 4144 | 173 | 40 |
| **Jag1** | Notch1 | 1811 | 2535 | 41 |
| **L1cam** | Itgav | 705 | 3688 | 42 |
| **Nlgn3** | Nrxn1 | 2037 | 2422 | 43 |
| **Hspg2** | Ptprs | 2912 | 1554 | 44 |
| **Shh** | Smo | 4144 | 446 | 45 |
| **Sema4a** | Plxnd1 | 2140 | 2573 | 46 |
| **Lama5** | Sdc1 | 1057 | 3661 | 47 |
| **Col4a1** | Itgb8 | 3909 | 1100 | 48 |
| **Sema6a** | Plxna2 | 92 | 5011 | 49 |
| **Lamc1** | Itgav | 1441 | 3688 | 50 |
| **Col4a1** | Itga1 | 3909 | 1225 | 51 |
| **Fgf18** | Fgfr1 | 4294 | 888 | 52 |
| **Hspg2** | Itgb1 | 2912 | 2286 | 53 |
| **Efna4** | Epha3 | 53 | 5167 | 54 |
| **Ihh** | Boc | 5074 | 173 | 55 |
| **Gm11808** | Egfr | 3949 | 1299 | 56 |
| **Fgf17** | Fgfr1 | 4395 | 888 | 57 |
| **Lamb1** | Itgav | 1668 | 3688 | 58 |
| **Sema3f** | Plxna3 | 1051 | 4444 | 59 |
| **Sema3f** | Plxna1 | 1051 | 4650 | 60 |
| **Ltbp1** | Itgb5 | 3136 | 2719 | 61 |
| **L1cam** | Itga5 | 705 | 5159 | 62 |
| **Efna4** | Epha7 | 53 | 5902 | 63 |
| **Ctf1** | Lifr | 379 | 5686 | 64 |
| **Bmp8b** | Bmpr1a | 3723 | 2345 | 65 |
| **Bmp8b** | Acvr1 | 3723 | 2361 | 66 |
| **Gdf5** | Ror2 | 5373 | 786 | 67 |
| **Mfap2** | Notch1 | 3648 | 2535 | 68 |
| **Col4a1** | Itgb1 | 3909 | 2286 | 69 |
| **Wnt5a** | Fzd1 | 5788 | 416 | 70 |
| **Vegfa** | Itga9 | 6167 | 130 | 71 |
| **Gm11808** | Acvr1 | 3949 | 2361 | 72 |
| **Lama1** | Itgb8 | 5217 | 1100 | 73 |
| **Jag1** | Notch4 | 1811 | 4507 | 74 |
| **Gm11808** | Agtr1a | 3949 | 2400 | 75 |
| **Wnt5a** | Ror1 | 5788 | 573 | 76 |
| **Vegfa** | Nrp1 | 6167 | 215 | 77 |
| **Bcan** | Egfr | 5100 | 1299 | 78 |
| **Lama1** | Itga1 | 5217 | 1225 | 79 |
| **Gm11808** | Notch1 | 3949 | 2535 | 80 |
| **Kitl** | Kit | 114 | 6397 | 81 |
| **Cdh1** | Cdh2 | 261 | 6264 | 82 |
| **Serpine2** | Lrp1 | 4321 | 2219 | 83 |
| **Hspg2** | Sdc1 | 2912 | 3661 | 84 |
| **Wnt5a** | Ror2 | 5788 | 786 | 85 |
| **App** | Lrp1 | 4431 | 2219 | 86 |
| **Myoc** | Fzd1 | 6289 | 416 | 87 |
| **Efna4** | Epha1 | 53 | 6700 | 88 |
| **Col18a1** | Itga5 | 1610 | 5159 | 89 |
| **Dusp18** | Itga1 | 5562 | 1225 | 90 |
| **Tgfa** | Egfr | 5582 | 1299 | 91 |
| **Timp2** | Itgb1 | 4792 | 2286 | 92 |
| **Il11** | Gm13305 | 6455 | 633 | 93 |
| **Tac1** | Tacr3 | 2816 | 4289 | 94 |
| **Efnb2** | Epha3 | 2019 | 5167 | 95 |
| **Bmp8b** | Acvr2a | 3723 | 3481 | 96 |
| **Edn1** | Ednra | 6997 | 208 | 97 |
| **Il7** | Il2rg | 3521 | 3697 | 98 |
| **Ncam1** | Fgfr1 | 6380 | 888 | 99 |
| **Dhh** | Boc | 7140 | 173 | 100 |

Supplemental Table 1 UB-SP3

| Ligand | Receptor | Ligand rank | Receptor rank | Interaction rank |
|---|---|---|---|---|
| **Slit2** | Robo1 | 139 | 311 | 1 |
| **Efna4** | Epha7 | 53 | 597 | 2 |
| **Slit2** | Robo2 | 139 | 537 | 3 |
| **Cdh1** | Egfr | 261 | 439 | 4 |
| **Btc** | Egfr | 322 | 439 | 5 |
| **L1cam** | Egfr | 705 | 439 | 6 |
| **Bmp7** | Bmpr1a | 59 | 1247 | 7 |
| **Npy** | Fap | 1017 | 363 | 8 |
| **Fgf9** | Fgfr1 | 1118 | 513 | 9 |
| **Cdh1** | Cdh2 | 261 | 1499 | 10 |
| **Pdgfc** | Pdgfra | 1664 | 424 | 11 |
| **Kitl** | Kit | 114 | 2151 | 12 |
| **Sema3c** | Plxnd1 | 771 | 1618 | 13 |
| **Sema3d** | Nrp1 | 749 | 1661 | 14 |
| **Sema3c** | Nrp1 | 771 | 1661 | 15 |
| **Fgf12** | Fgfr1 | 1951 | 513 | 16 |
| **Kitl** | Epor | 114 | 2414 | 17 |
| **Bmp7** | Eng | 59 | 2586 | 18 |
| **Sema3f** | Nrp1 | 1051 | 1661 | 19 |
| **L1cam** | Ephb2 | 705 | 2153 | 20 |
| **Pdgfc** | Pdgfrb | 1664 | 1304 | 21 |
| **Slit2** | Sdc1 | 139 | 3092 | 22 |
| **Bmp7** | Acvr1 | 59 | 3234 | 23 |
| **Cdh1** | Itgae | 261 | 3048 | 24 |
| **Sema4a** | Plxnd1 | 2140 | 1618 | 25 |
| **Pdgfa** | Pdgfra | 3390 | 424 | 26 |
| **Nrtn** | Gfra1 | 17 | 3947 | 27 |
| **Col4a1** | Itgb8 | 3909 | 113 | 28 |
| **Lama5** | Sdc1 | 1057 | 3092 | 29 |
| **Efnb2** | Ephb2 | 2019 | 2153 | 30 |
| **Sema3f** | Plxna1 | 1051 | 3162 | 31 |
| **Shh** | Boc | 4144 | 105 | 32 |
| **Gm11808** | Egfr | 3949 | 439 | 33 |
| **Sema3f** | Plxna3 | 1051 | 3409 | 34 |
| **Tac1** | Tacr3 | 2816 | 1835 | 35 |
| **Pdgfa** | Pdgfrb | 3390 | 1304 | 36 |
| **Hspg2** | Ptprs | 2912 | 1812 | 37 |
| **Prss23** | Tmem222 | 1609 | 3148 | 38 |
| **Wnt7b** | Fzd1 | 3770 | 1024 | 39 |
| **Fgf18** | Fgfr1 | 4294 | 513 | 40 |
| **Cgn** | F11r | 2929 | 1947 | 41 |
| **Fgf17** | Fgfr1 | 4395 | 513 | 42 |
| **Bmp8b** | Bmpr1a | 3723 | 1247 | 43 |
| **Shh** | Smo | 4144 | 891 | 44 |
| **Ihh** | Boc | 5074 | 105 | 45 |
| **Bmp7** | Bmpr2 | 59 | 5145 | 46 |
| **Lamc2** | Itgb1 | 172 | 5139 | 47 |
| **Cntn2** | Nrp1 | 3656 | 1661 | 48 |
| **Lama1** | Itgb8 | 5217 | 113 | 49 |
| **L1cam** | Itga5 | 705 | 4734 | 50 |
| **Nlgn3** | Nrxn1 | 2037 | 3410 | 51 |
| **Npnt** | Itgb1 | 338 | 5139 | 52 |
| **Bcan** | Egfr | 5100 | 439 | 53 |
| **Cdh1** | Igf1r | 261 | 5306 | 54 |
| **Btc** | Erbb4 | 322 | 5315 | 55 |
| **Cdh1** | Lrp5 | 261 | 5432 | 56 |
| **Shh** | Cdon | 4144 | 1550 | 57 |
| **Efnb2** | Ephb3 | 2019 | 3684 | 58 |
| **Efna5** | Epha7 | 5351 | 597 | 59 |
| **Wnt5a** | Ror1 | 5788 | 161 | 60 |
| **Efnb2** | Ephb4 | 2019 | 3962 | 61 |
| **Hspg2** | Sdc1 | 2912 | 3092 | 62 |
| **Tgfa** | Egfr | 5582 | 439 | 63 |
| **Lama5** | Itgb1 | 1057 | 5139 | 64 |
| **Jag1** | Notch2 | 1811 | 4420 | 65 |
| **Eda** | Edar | 4171 | 2083 | 66 |
| **Nrtn** | Gfra2 | 17 | 6297 | 67 |
| **Col18a1** | Itga5 | 1610 | 4734 | 68 |
| **Efna1** | Epha7 | 5758 | 597 | 69 |
| **Vegfa** | Itga9 | 6167 | 245 | 70 |
| **Efna4** | Epha5 | 53 | 6383 | 71 |
| **Gdf5** | Ror2 | 5373 | 1064 | 72 |
| **Lamc1** | Itgb1 | 1441 | 5139 | 73 |
| **Wnt11** | Fzd7 | 41 | 6546 | 74 |
| **Vegfa** | Egfr | 6167 | 439 | 75 |
| **Gdf5** | Bmpr1a | 5373 | 1247 | 76 |
| **Ihh** | Cdon | 5074 | 1550 | 77 |
| **Il11** | Gm13305 | 6455 | 193 | 78 |
| **Ncam1** | Robo1 | 6380 | 311 | 79 |
| **Anxa1** | Egfr | 6255 | 439 | 80 |
| **Col18a1** | Itgb1 | 1610 | 5139 | 81 |
| **Lamb1** | Itgb1 | 1668 | 5139 | 82 |
| **Wnt5a** | Fzd1 | 5788 | 1024 | 83 |
| **Wnt5a** | Ror2 | 5788 | 1064 | 84 |
| **Ncam1** | Fgfr1 | 6380 | 513 | 85 |
| **Jag1** | Notch1 | 1811 | 5097 | 86 |
| **Wnt5a** | Fzd2 | 5788 | 1160 | 87 |
| **Bmp8b** | Acvr1 | 3723 | 3234 | 88 |
| **Fgf16** | Fgfr1 | 6627 | 513 | 89 |
| **Gm11808** | Acvr1 | 3949 | 3234 | 90 |
| **Dhh** | Boc | 7140 | 105 | 91 |
| **Itgb3bp** | Itgb5 | 1539 | 5717 | 92 |
| **L1cam** | Itgav | 705 | 6560 | 93 |
| **Epgn** | Egfr | 6873 | 439 | 94 |
| **Myoc** | Fzd1 | 6289 | 1024 | 95 |
| **Wnt4** | Fzd2 | 6163 | 1160 | 96 |
| **Grp** | Grpr | 3325 | 4033 | 97 |
| **Efna5** | Ephb2 | 5351 | 2153 | 98 |
| **Bmp3** | Bmpr1a | 6286 | 1247 | 99 |
| **Jag1** | Notch4 | 1811 | 5833 | 100 |

Supplemental Table 1 UB-RV

| Ligand | Receptor | Ligand rank | Receptor rank | Interaction rank |
|---|---|---|---|---|
| **Efna4** | Epha4 | 53 | 113 | 1 |
| **Bmp7** | Acvr2b | 59 | 281 | 2 |
| **Slit2** | Sdc1 | 139 | 329 | 3 |
| **Cdh1** | Cdh2 | 261 | 438 | 4 |
| **Slit2** | Gpc1 | 139 | 712 | 5 |
| **Efna4** | Epha7 | 53 | 1044 | 6 |
| **Fgf9** | Fgfr1 | 1118 | 26 | 7 |
| **Lama5** | Sdc1 | 1057 | 329 | 8 |
| **Lamc2** | Itga6 | 172 | 1394 | 9 |
| **L1cam** | Fgfr2 | 705 | 1003 | 10 |
| **Jag1** | Notch1 | 1811 | 157 | 11 |
| **Fgf12** | Fgfr1 | 1951 | 26 | 12 |
| **Fgf9** | Fgfr2 | 1118 | 1003 | 13 |
| **Efnb2** | Epha4 | 2019 | 113 | 14 |
| **Col18a1** | Gpc1 | 1610 | 712 | 15 |
| **Slit2** | Robo1 | 139 | 2214 | 16 |
| **Lama5** | Itga6 | 1057 | 1394 | 17 |
| **Lamc2** | Cd151 | 172 | 2323 | 18 |
| **Lama5** | Bcam | 1057 | 1452 | 19 |
| **Lamc2** | Itga3 | 172 | 2445 | 20 |
| **Cdh1** | Ptprm | 261 | 2502 | 21 |
| **Lamc1** | Itga6 | 1441 | 1394 | 22 |
| **Fgf12** | Fgfr2 | 1951 | 1003 | 23 |
| **Cdh1** | Igf1r | 261 | 2769 | 24 |
| **Lamb1** | Itga6 | 1668 | 1394 | 25 |
| **Hspg2** | Col13a1 | 2912 | 158 | 26 |
| **Hspg2** | Sdc1 | 2912 | 329 | 27 |
| **Wnt11** | Fzd7 | 41 | 3385 | 28 |
| **Jag1** | Notch2 | 1811 | 1618 | 29 |
| **Lama5** | Itga3 | 1057 | 2445 | 30 |
| **Nrtn** | Gfra1 | 17 | 3510 | 31 |
| **Efnb2** | Ephb1 | 2019 | 1571 | 32 |
| **Bmp7** | Bmpr1a | 59 | 3658 | 33 |
| **Cdh1** | Ptprf | 261 | 3463 | 34 |
| **Mfap2** | Notch1 | 3648 | 157 | 35 |
| **Lamc1** | Itga3 | 1441 | 2445 | 36 |
| **Lamc2** | Itgb4 | 172 | 3720 | 37 |
| **Bmp8b** | Acvr2b | 3723 | 281 | 38 |
| **Gm11808** | Notch1 | 3949 | 157 | 39 |
| **Lamb1** | Itga3 | 1668 | 2445 | 40 |
| **Nlgn3** | Nrxn2 | 2037 | 2100 | 41 |
| **Wnt7b** | Fzd10 | 3770 | 486 | 42 |
| **Slit2** | Robo2 | 139 | 4134 | 43 |
| **Fgf18** | Fgfr1 | 4294 | 26 | 44 |
| **Fgf17** | Fgfr1 | 4395 | 26 | 45 |
| **App** | Ngfr | 4431 | 175 | 46 |
| **Lama5** | Itgb4 | 1057 | 3720 | 47 |
| **Spint1** | St14 | 368 | 4549 | 48 |
| **App** | Gpc1 | 4431 | 712 | 49 |
| **Lamc1** | Itgb4 | 1441 | 3720 | 50 |
| **Agrn** | Lrp4 | 3593 | 1693 | 51 |
| **Fgf18** | Fgfr2 | 4294 | 1003 | 52 |
| **Jag1** | Notch4 | 1811 | 3487 | 53 |
| **Hspg2** | Ptprs | 2912 | 2473 | 54 |
| **Lamb1** | Itgb4 | 1668 | 3720 | 55 |
| **Fgf17** | Fgfr2 | 4395 | 1003 | 56 |
| **Efna5** | Epha4 | 5351 | 113 | 57 |
| **Prss23** | Tmem222 | 1609 | 3861 | 58 |
| **Cgn** | Tgfbr1 | 2929 | 2619 | 59 |
| **Gdf5** | Acvr2b | 5373 | 281 | 60 |
| **Cd24a** | Selp | 105 | 5619 | 61 |
| **Oxt** | Oxtr | 5503 | 293 | 62 |
| **Efna1** | Epha4 | 5758 | 113 | 63 |
| **Sema4d** | Plxnb1 | 4111 | 1781 | 64 |
| **Lama1** | Gpc1 | 5217 | 712 | 65 |
| **Fgf9** | Fgfr4 | 1118 | 4853 | 66 |
| **L1cam** | Ephb2 | 705 | 5347 | 67 |
| **Inhba** | Acvr2b | 6107 | 281 | 68 |
| **Sema3c** | Plxnd1 | 771 | 5617 | 69 |
| **Efna5** | Epha7 | 5351 | 1044 | 70 |
| **Ncam1** | Fgfr1 | 6380 | 26 | 71 |
| **Serpinc1** | Gpc1 | 5752 | 712 | 72 |
| **Cgn** | Ocln | 2929 | 3615 | 73 |
| **Bmp3** | Acvr2b | 6286 | 281 | 74 |
| **Gm11808** | Tgfbr1 | 3949 | 2619 | 75 |
| **Inhba** | Bambi | 6107 | 499 | 76 |
| **Lama1** | Itga6 | 5217 | 1394 | 77 |
| **Fgf16** | Fgfr1 | 6627 | 26 | 78 |
| **Myoc** | Fzd3 | 6289 | 380 | 79 |
| **Sema3c** | Nrp2 | 771 | 5933 | 80 |
| **Myoc** | Fzd10 | 6289 | 486 | 81 |
| **Efna1** | Epha7 | 5758 | 1044 | 82 |
| **Fgf12** | Fgfr4 | 1951 | 4853 | 83 |
| **Lamb3** | Itga6 | 5432 | 1394 | 84 |
| **Vegfa** | Gpc1 | 6167 | 712 | 85 |
| **Efna5** | Ephb1 | 5351 | 1571 | 86 |
| **Il2** | Ngfr | 6756 | 175 | 87 |
| **Dusp18** | Itga6 | 5562 | 1394 | 88 |
| **Hhipl2** | Cachd1 | 5382 | 1599 | 89 |
| **Sema3f** | Nrp2 | 1051 | 5933 | 90 |
| **Col18a1** | Gpc4 | 1610 | 5404 | 91 |
| **App** | Slc45a3 | 4431 | 2593 | 92 |
| **Rspo4** | Lgr4 | 5392 | 1708 | 93 |
| **Pigf** | Flt1 | 341 | 6883 | 94 |
| **Timp2** | Itga3 | 4792 | 2445 | 95 |
| **Ntf3** | Ngfr | 7067 | 175 | 96 |
| **Sema3f** | Plxna3 | 1051 | 6193 | 97 |
| **Kitl** | Epor | 114 | 7140 | 98 |
| **Tac1** | Tacr1 | 2816 | 4511 | 99 |
| **Efna1** | Ephb1 | 5758 | 1571 | 100 |
